# Top-down modulation of the retinal code via histaminergic neurons of the hypothalamus

**DOI:** 10.1101/2022.04.26.489509

**Authors:** Rebekah A. Warwick, Serena Riccitelli, Alina S. Heukamp, Hadar Yaakov, Lea Ankri, Jonathan Mayzel, Noa Gilead, Reut Parness-Yossifon, Michal Rivlin-Etzion

## Abstract

The mammalian retina is considered an autonomous circuit, yet work dating back to Ramon y Cajal indicates that it receives inputs from the brain. How such inputs affect retinal processing has remained unknown. We identified brain-to-retina projections of histaminergic neurons from the mouse hypothalamus, which densely innervated the dorsal retina. Histamine application, or chemogenetic activation of histaminergic axons, altered spontaneous and light-evoked activity of various retinal ganglion cells (RGCs), including direction-selective RGCs. These cells exhibited broader directional tuning and gained responses to high motion velocities. Such changes could improve vision when objects move fast across the visual field (e.g. while running), which fits with the known increased activity of histaminergic neurons during arousal. In humans, an antihistamine drug non-uniformly modulated visual sensitivity across the visual field, indicating an evolutionary conserved function of the histaminergic system. Our findings expose a previously unappreciated role for brain-to-retina projections in modulating retinal function.

## Introduction

The retina is typically viewed as an autonomous neuronal tissue, which processes external input – the visual image – and projects its output to the brain. Yet, more than a century ago, Ramon y Cajal showed that the avian retina is innervated by retinopetal axons coming from the brain via the optic nerve (Ramón y Cajal, 1888, 1889), suggesting that visual processing in the retina is subject to top-down modulations. Later, the presence of retinopetal axons was confirmed in various other vertebrate species, with some reports also in mammals, including humans (reviewed in Repérant et al. (1989)). These studies described a few fibers that emerged from the optic disc and branched extensively to cover a large portion of the retina and tended to terminate in the inner plexiform layer or the inner nuclear layer (Dräger et al., 1984; Perry et al., 1984). Still, retinopetal axons in the mammalian retina remain elusive and their origin is controversial, probably due to the small number of projecting neurons.

Immunohistochemical analyses demonstrated the presence of retinopetal axons containing histamine in guinea pig, mouse, rat and primate retinas (Airaksinen and Panula, 1988; Gastinger et al., 1999, 2001; Greferath et al., 2009; Koves and Csaki, 2016). Since neurons located in the tuberomammillary nucleus (TMN) of the posterior hypothalamus are the only source of neuronal histamine in the mammalian nervous system (Airaksinen and Panula, 1988; Manning et al., 1996; Panula et al., 1984; Watanabe et al., 1984), it was suggested that histaminergic retinopetal axons originate from the TMN (Gastinger et al., 2006a). Moreover, it was explicitly shown that the retina does not contain any histamine-producing neurons (Gastinger et al., 2006a), yet retinal histamine levels are comparable with other brain regions innervated by histaminergic neurons (Arbonés et al., 1988; Nowak, 1990). Histaminergic axons generally do not form synaptic contacts, so histamine is thought to act in a paracrine fashion (Greferath et al., 2009; Haas and Panula, 2003) via three types of histamine receptors (HRs) that have been identified in the mammalian CNS. H_1_R and H_2_R are G_q_- and G_s_-coupled receptors, respectively, and their direct action is usually excitatory, whereas H_3_R is a G_i_-coupled receptor that typically has an inhibitory effect (Haas and Panula, 2003). All three HRs were found in the retina (Arbonés et al., 1988; Nowak, 1990), suggesting that histamine plays a neuromodulatory role in this tissue (Frazão et al., 2011; Gastinger et al., 2006b; Greferath et al., 2009; Horio et al., 2018; Vila et al., 2012; Yu et al., 2011, 2009).

Histaminergic retinopetal axons pose a potential paradigm shift, as their existence suggests that higher brain areas can shape the retinal code. However, the functional role of histamine and its contribution to early visual processing is still poorly understood. Previous studies demonstrated that histamine acts on several retinal cell types, including cones, bipolar cells and amacrine cells, via the activation of different receptors (Frazão et al., 2011; Horio et al., 2018; Vila et al., 2012; Yu et al., 2009). Two studies found that histamine alters the output of a large portion of retinal ganglion cells (RGCs), but this effect was highly variable (Akimov et al., 2010; Gastinger et al., 2004).

The firing rate of histaminergic neurons is correlated with the arousal state of the animal; they are minimally active during sleep and their activity peaks during attentive waking (Haas et al., 2008; Scammell et al., 2019; Takahashi et al., 2006; Vanni-Mercier et al., 2003). It has recently been suggested that arousal state directly influences activity in early visual structures – the dorsal lateral geniculate nucleus (dLGN) and the superior colliculus (SC) (Liang et al., 2020; Schröder et al., 2020), but it is unknown whether these effects are mediated by histamine or even retinopetal axons. To further complicate matters, histaminergic neurons are known to project to many brain areas, including those responsible for visual processing, raising the possibility that they locally modulate RGC axon terminals in these regions (Gastinger et al., 2006a; Uhlrich et al., 2002; Yu et al., 2015).

Here, we sought to reveal the effects of histaminergic retinopetal projections on retinal output. We first used viral injections in transgenic mice and identified histaminergic retinopetal projections which originate in the TMN. Using two-photon Ca^2+^ imaging, multielectrode array (MEA) and targeted patch clamp recordings, we showed that histamine affects both the spontaneous and light-evoked activity of various RGCs, including the OFF-transient alpha RGC and direction-selective ganglion cell (DSGC). Crucially, we demonstrated that the selective chemogenetic activation of the histaminergic retinopetal axons induces significant changes in RGCs’ activity, including that of DSGCs. In DSGCs, histamine broadened directional tunings and enhanced the responses to higher motion velocities. Finally, we found that an antihistamine can affect visual sensitivity non-uniformly across the visual field in human subjects.

## Results

### The retina is innervated by histaminergic fibers arising from the tuberomammillary nucleus of the hypothalamus

To confirm the presence of retinopetal axons and their origin, we unilaterally injected an adeno-associated virus (AAV) encoding for a fluorescent marker into the TMN of the posterior hypothalamus of wild-type mice (Figure S1A). After 4-6 weeks, we observed fluorescence in cell bodies in the TMN area. Immunostaining showed that some of these neurons were also positive for a specific marker for histaminergic cells, histidine decarboxylase (HDC), the enzyme that catalyzes the final step in the synthesis of histamine (Figure S1B). In addition to the labeled cell somas, we detected labeled axons in all regions known to be targeted by the TMN, including the retina, where few axons emerged from the optic disc and predominantly innervated the dorsal part of the retina (Figure S1C). Using this approach, we also observed labeled cell bodies in the ganglion cell layer of the retina, which possibly originated from retrograde labeling of RGC axons passing close to the injection site (Figure S1C). Therefore, to ensure we specifically target histaminergic neurons, we switched to using HDC-Cre mice.

The HDC-Cre mice experiments entailed injecting a Cre-dependent AAV encoding for a fluorescent marker into the TMN (Figure 1A). Fluorescent cells were restricted to the TMN area and immunoreactive to HDC (Figure 1B), validating the specificity of Cre-mediated recombination in HDC^+^ cells. Optic nerve wholemounts confirmed the presence of a few centrifugal fibers running through the optic nerve (Figure 1C, red arrowheads). We also observed axons that seem to terminate in the optic nerve (Figure 1C, red dashed line), but we cannot exclude the option that they reach the retina but are not detected by our approach. Notably, we found strongly labeled retinopetal axons in the retina that ran from the optic disc and covered its dorsal half (Figure 1D). The specificity of the virus allowed us to undoubtedly identify these retinopetal axons as histaminergic. In addition, it prevented the appearance of labeled neurons in the retina and allowed us to localize the histaminergic axons in the retinal layers. Typically, these axons ran in the nerve fibers layer (NFL) and ganglion cell layer (GCL) and descended orthogonally into the inner plexiform layer (IPL), where, despite their small number, they branched extensively and covered a large portion of the retina, reaching the ora serrata (Figures 1D and 1E). These results are in agreement with previous anatomical studies (Gastinger et al., 2006a; Greferath et al., 2009). The existence of histaminergic retinopetal projections suggests that retinal neurons can be subject to top-down neuromodulatory influences that act on the early stages of visual processing.

**Figure 1.**
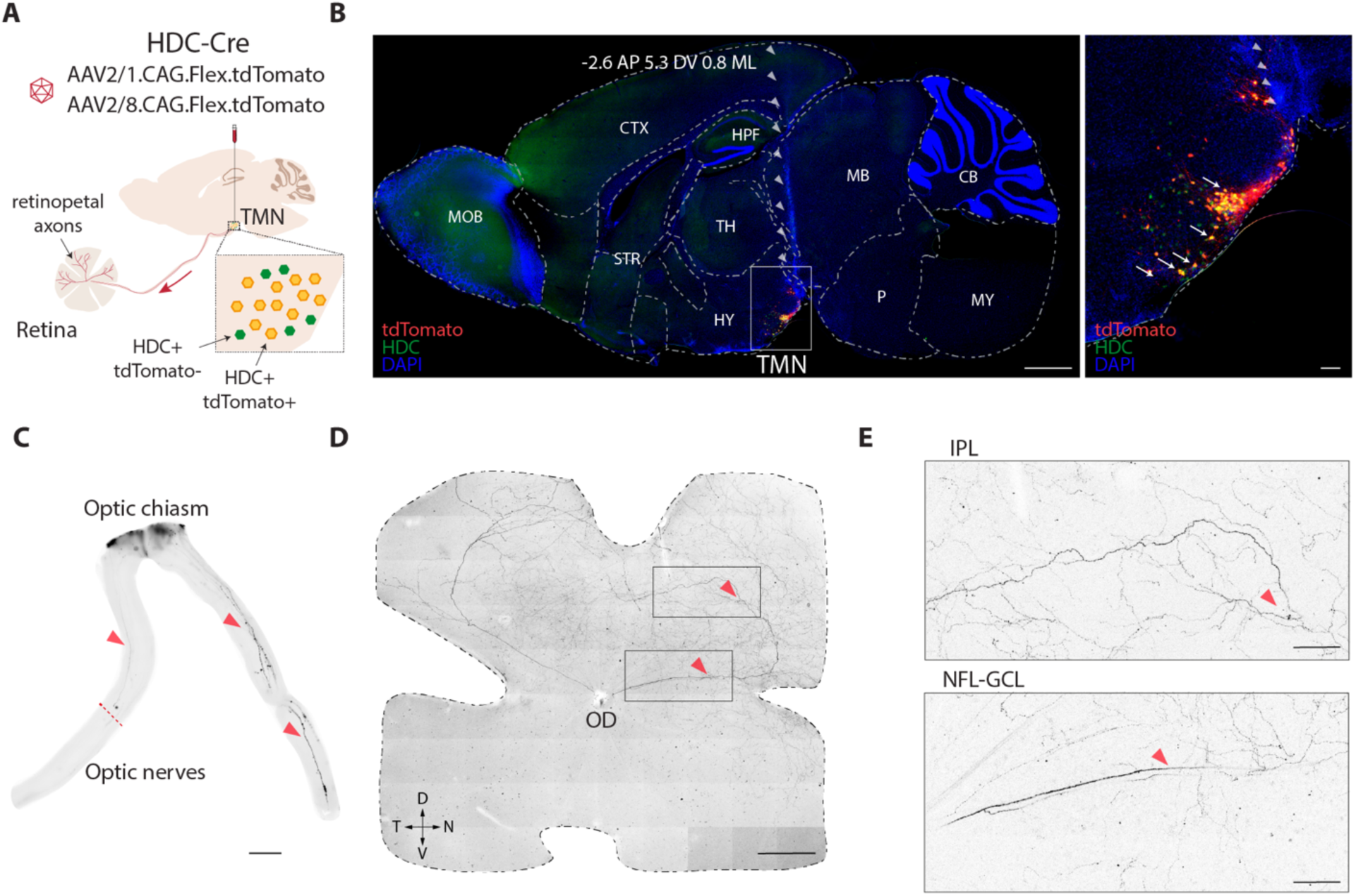
Retinopetal histaminergic projections in HDC-Cre mice. (**A**) Microinjection of viral tracers in the TMN of transgenic mice expressing Cre in HDC^+^ cells. Brains and retinas were analyzed after 4-6 weeks. (**B**) Left: sagittal brain slice of an HDC-Cre mouse 4 weeks after injection of AAV2/8.CAG.Flex.tdTomato showing immunohistochemistry of HDC^+^ (green) and tdTomato^+^ (red) cells. Gray arrowheads indicate the injection site. Right: High-magnification of the region indicated by the white box showing double-labeled cell bodies (white arrows). Scale bars, 1000 µm, left panel; 100 µm, right panel. (**C**) Tiled fluorescence images showing histaminergic retinopetal axons in wholemount optic nerves. Red arrowheads indicate two positively labeled histaminergic fibers emerging from the optic chiasm (right). On the left, a dashed line indicates a fiber that terminates in the optic nerve. Scale bar, 500 µm. (**D**) Tiled fluorescence image of the retina (maximum intensity projection) showing two major fibers in the dorsal retina. The primary axons emerge from the optic disc. Scale bar, 500 µm. (**E**) Higher magnification of regions indicated by black boxes in (D). Top: Histaminergic axons branching in the IPL. Bottom: Histaminergic axon branches run through the NFL-GCL. Scale bars, 100 µm. Abbreviations: CTX – Cortex, HPF – Hippocampal formation, TH – Thalamus, HY – Hypothalamus, TMN – Tuberomammillary nucleus, CB – Cerebellum, MB – Midbrain, P – Pons, MY – Medulla, IPL – Inner plexiform layer, GCL – Ganglion cell layer, NFL – Nerve fiber layer, OD – Optic disc, D, N, T, V – dorsal, nasal, temporal, ventral.

### Histamine drives changes in retinal ganglion cells’ firing rates

Retinal research is generally carried out *ex vivo*, where inputs from the brain to the retina are severed. Under these conditions, retinopetal axons are no longer active, thus preventing the assessment of any histamine modulatory effect that might occur *in vivo*. Therefore, we opted to use multi-electrode array (MEA) recordings to establish the dose-response (1-20 µM) effect of histamine bath application on the baseline firing rate of spiking neurons in the RGC layer. We refer to these neurons as RGCs, although displaced spiking amacrine cells can also be recorded (Greschner et al., 2014). The histamine dose was based on previous functional experiments of the retina that used between 0.5-100 µM histamine (Akimov et al., 2010; Frazão et al., 2011; Gastinger et al., 2004; Horio et al., 2018; Vila et al., 2012; Yu et al., 2009). We found that 2 µM histamine was the lowest concentration that caused significant increases in the baseline firing rates; at 5 µM, the change in firing rate and percentage of responsive RGCs reached a plateau (Figure S2). Therefore, for further experiments, we used a histamine concentration in the range of 5-20 µM.

Next, we sought to establish the fraction of RGCs that respond to histamine. Bath application of 5 µM histamine to isolated retinas on a MEA caused some RGCs to sharply increase their basal firing rate, while others were not affected (Figure 2A). In the control experiments, histamine was not added. In total, histamine increased the firing rate in 46.0±13.9% (mean±SD) of RGCs (6 retinas), significantly more than in the control data set (7 retinas) (Figures 2B and 2C). Only 1.1±1.1% of RGCs showed reduced spiking activity upon histamine application. RGCs responsive to histamine included a variety of subtypes with ON, OFF, and ON-OFF polarity preferences, characterized by their response to a full-field stimulus given prior to the addition of histamine (Figure 2D).

**Figure 2.**
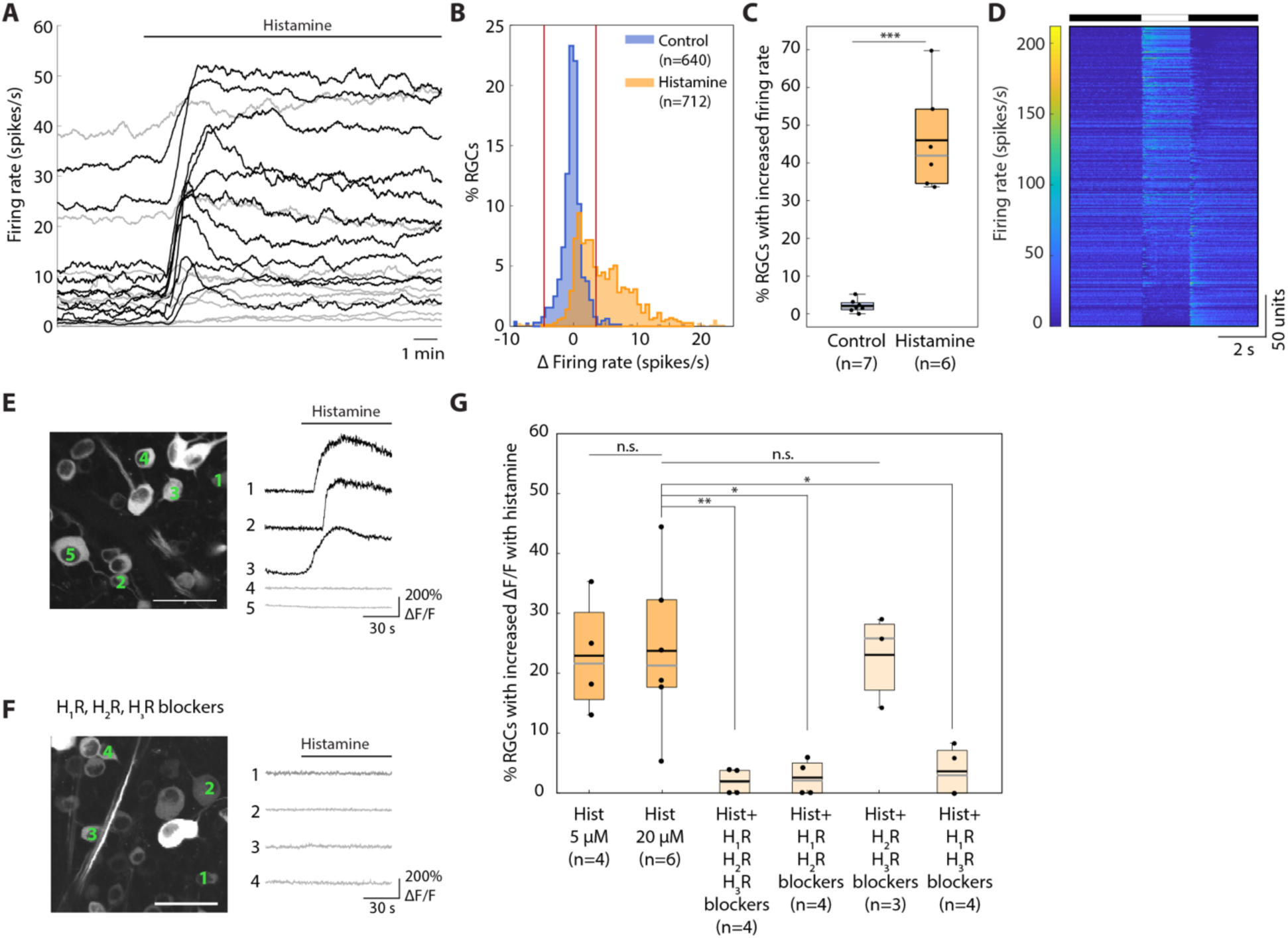
Histamine increases the baseline activity of RGCs. (**A**) Baseline firing rates of example RGCs recorded on the MEA with bath application of histamine. Black cells significantly increased their firing rate upon the addition of 5 µM histamine application (horizontal black line above), while gray cells did not. (**B**) Distribution of the difference in firing rate compared to the baseline for control (no histamine added, blue) and histamine (orange) experiments. Red vertical lines show upper and lower 2.5% quantile of control distribution (−4.6 and 3.5 spikes/s, respectively). Distributions are significantly different (*p*«0.0001, Kolmogorov-Smirnov test). (**C**) The proportion of RGCs that significantly increased their firing rate is significantly greater with histamine (orange, n=6 retinas) than in the control (blue, n=7 retinas, *p*=0.0003, one-tailed two-sample t-test for unequal variances). Black and gray lines show the mean and median, respectively. (**D**) Responses to a full-field stimulus (indicated above) of all RGCs that increased their firing rate with histamine, sorted according to their ON-OFF preference, showing that various RGC types are responsive to histamine (n=363 RGCs). (**E**) Left: Two-photon image of neurons in the ganglion cell layer of a Thy1-GCamp6f mouse. Scale bar, 50 µm. Right: Ca^2+^ traces of three RGCs that responded to histamine application (20 µM, black) and two RGCs that did not (gray). (**F**) Same as in (E) but in the presence of blockers of H_1_R (Cetirizine, 20 µM), H_2_R (Famotidine, 40 µM) and H_3_R (JNJ 5207852, 20 µM). (**G**) Percentage of RGCs per retina that reacted to histamine (ΔF/F increased by more than 6 SDs relative to the baseline) when added alone (5 and 20 µM) or when histamine (20 µM) was added in the presence of various histamine receptor blocker combinations: H_1_R+H_2_R+H_3_R blockers – p=0.0099, H_1_R+H_2_R blockers – p=0.0107, H_1_R+H_3_R blockers – p=0.0120, one-way Welch’s ANOVA. Black and gray lines show the mean and median, respectively. *: p<0.05, **: *p*<0.01, ***: *p*<0.0001.

Consistently, experiments using two-photon Ca^2+^ imaging of neurons in the RGC layer expressing GCaMP6f (referred to as RGCs hereafter) showed that histamine increased the intracellular Ca^2+^ concentration in 22.9±9.6% and 23.7±13.5% of RGCs (mean±SD, 4 and 6 retinas) for 5 µM and 20 µM, respectively (Figures 2E and 2G). Blocking H_2_R and H_3_R did not change the percentage of RGCs that reacted to histamine, but any cocktail that included a H_1_R blocker significantly reduced the percentage of responsive RGCs, showing that the effect of histamine on baseline activity of RGCs is mainly mediated by the H_1_R (Figures 2F and 2G).

### Histamine alters RGC activity in both dorsal and ventral parts of the retina

Given that traced histaminergic retinopetal axons predominantly innervate the dorsal retina, we sought to understand whether histamine uniformly affects the output of RGCs located either in the dorsal or ventral areas of the retina. By separating the data obtained from bath application of histamine (Figure 2) according to retinal location, we found that both dorsal and ventral RGCs were responsive to histamine application (Figure S3). In the MEA recordings, 40.5±9.6% and 57.0±18.0% (mean±SD) of dorsal and ventral RGCs, respectively, significantly increased their baseline activity (Figures S3A and S3B). Similar results were obtained in the two-photon Ca^2+^ imaging experiments: 18.4±13.5% and 29.0±13.6% of dorsal and ventral RGCs, respectively, increased their intracellular Ca^2+^ concentration (Figure S3C). In order to understand how histamine may affect neurons in the ventral retina, we investigated the expression of H_1_R. We found H_1_R positive signal in all four retinal quadrants in both GCL and INL focal planes (Figure S3D).

### Histamine changes the light responses of RGCs

As histamine affects the baseline activity of RGCs, we investigated whether histamine modulates RGC responses to visual stimuli. Using two-photon Ca^2+^ imaging of the RGC layer, we characterized light responses of RGCs to a UV spot centered on the field of view (Figure 3A). RGCs were classified as either ON, OFF or non-responsive based on their Ca^2+^ transients (see *Methods*). Control experiments were undertaken in which histamine was not added. As the responses of some RGCs changed with time, only RGCs that kept their polarity preference (ON, OFF or non-responsive) in the pre and washout conditions were included. The light response of the majority of RGCs remained stable upon histamine application (20 µM). However, 27.5% of RGCs either lost, gained or changed their polarity to the spot stimulus (76/276 RGCs, 22 retinas) (Figures 3A and 3B; Figures S4A-C) compared to only 3% in the control data set (3/98 RGCs, 7 retinas; Figure 3B). Moreover, some of the RGCs that did not change their polarity preference exhibited an increase or decrease in their response amplitude (Figures 3A and 3C). The light-evoked response amplitude (calculated in units of standard deviation) before histamine application was plotted against the response amplitude after histamine application (Figure 3C). Deviation from the unity line was significantly larger in the histamine data set than in the control dataset (4.33±5.1 and 2.5±2.2 (mean±SD), respectively). These results suggest that histamine affects both the polarity preference and the light response amplitude of RGCs. Experiments were also carried out using 5 µM histamine. Under these conditions, histamine affected the light responses of a smaller fraction of RGCs (17.5% or 11/63, 5 retinas; Figures S4D and S4E). Nevertheless, the proportion of affected RGCs was significantly greater than that in the control data set.

**Figure 3.**
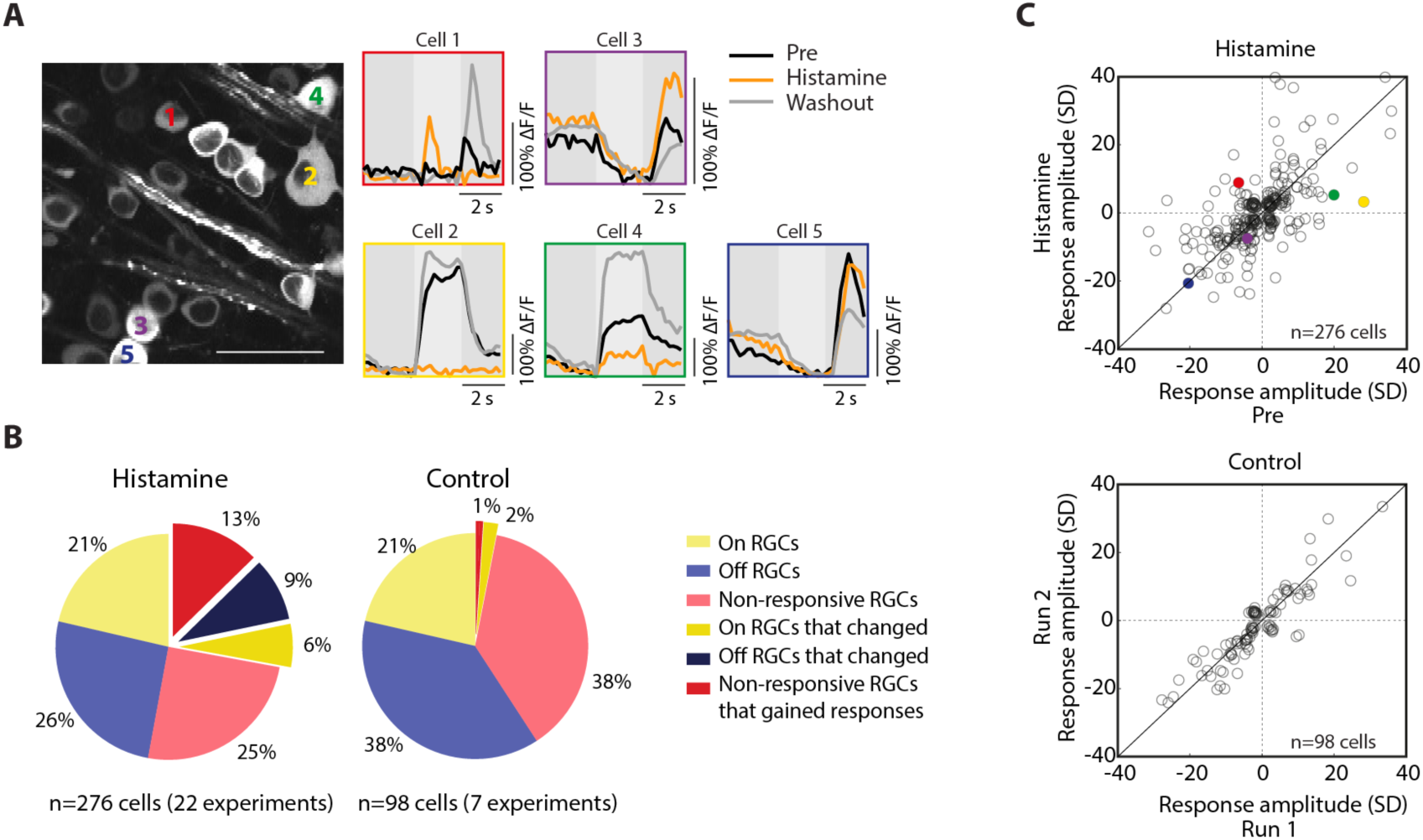
Two-photon Ca^2+^ imaging reveals that histamine affects light responses of RGCs to a spot stimulus. (**A**) Example experiment showing how histamine affects RGCs light responses to a UV spot stimulus. Left: Two-photon image of neurons in the ganglion cell layer. Scale bar, 50 µm. Right: Ca^2+^ traces of 5 RGCs before histamine (black), with histamine (orange) and after histamine washout (gray). (**B**) Pie charts showing the percentages of RGCs that changed their polarity preference in the histamine and control data sets (*p*<0.0001, Fisher’s exact test). (**C**) Population data showing how response amplitudes to a spot stimulus change with histamine (top) and in control experiments (bottom). Responses of ON RGCs (ON-OFF index > 0) are plotted as positive amplitudes, and those of OFF RGCs (ON-OFF index < 0) as negative response amplitudes. The colored dots in the upper plot correspond to the RGCs shown in (A). Deviation from the unity line is significantly larger in the histamine data set compared to the control data set *p*= 0.0045, Kolmogorov-Smirnov test.

Given that we found that dorsal and ventral RGCs are equally responsive to histamine, we next addressed whether histamine differentially affects their light responses. In the ventral retina, 22.4% (30/134, 10 retinas) of RGCs changed their polarity preference compared to 32.4% (46/142, 12 retinas) in the dorsal retina (Figure S5A). Histamine also had comparable effects on RGCs’ response amplitudes in the ventral and dorsal areas (Figure S5B).

The Ca^2+^ imaging data revealed a variety of histamine effects on the light-evoked responses of different RGCs. To understand how histamine influences the encoding properties of specific RGC subtypes, we conducted cell-attached recordings. We recorded from three alpha RGC subtypes (ON-sustained, OFF-sustained and OFF-transient) and from ON-OFF posterior preferring direction-selective ganglion cells (pDSGCs). Neither ON-sustained nor OFF-sustained alpha RGCs were affected by the application of histamine (20 µM). These cells’ baseline firing rates (Figures 4A_1-2_ and 4B_1-2_) and light responses to spot stimuli (Figure 4C_1-2_) remained unaltered by histamine. Conversely, OFF-transient alpha RGCs did increase their background firing rates after histamine was added (Figures 4A_3_ and 4B_3_). This change in background firing made inhibition at light onset more apparent (Figure 4C_3_). In addition, the maximum firing rate was significantly reduced for OFF-transient alpha RGCs when presented with larger spots (Figure 4D_3_), as were their response durations for smaller spots (Figure 4E_3_). Similarly, pDSGCs visibly increased their basal firing rate upon histamine application (Figure 4A_4_), with 3/7 RGCs entering into a depolarization block. This increase was also observable in their background firing rates during spot stimuli (Figure 4B_4_). The pDSCGs’ light responses to spots were poor compared to those of the alpha RGC subtypes (Figures 4C-E) and tended to be further diminished by histamine (Figure 4C_4_). However, this reduction was only significant for the OFF response duration (Figure 4E_4_).

**Figure 4.**
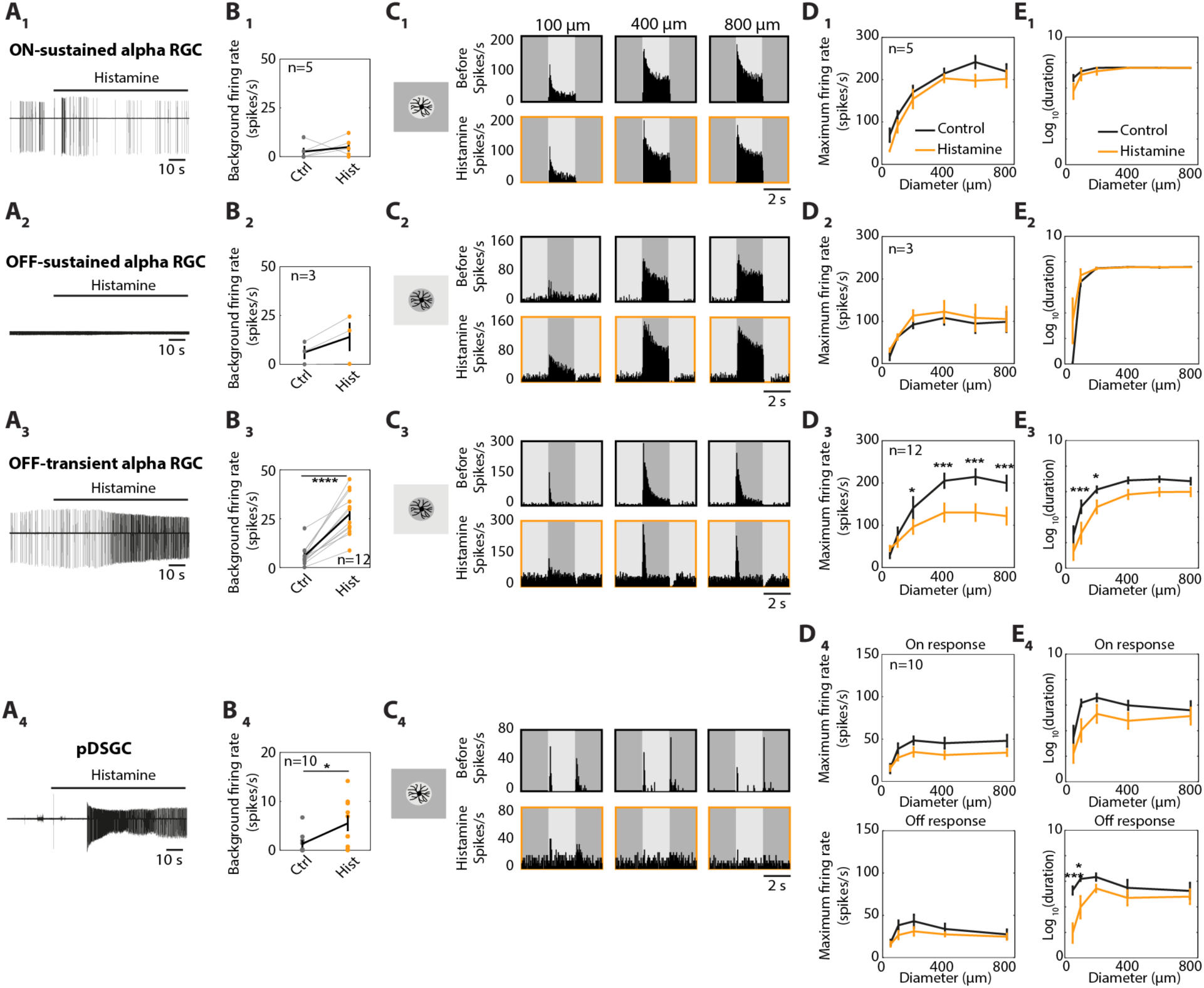
Histamine changes the spontaneous activity and light responses of specific RGC subtypes. (**A**) Example traces (cell-attached recordings) of spiking activity of an ON-sustained alpha RGC (A_1_), OFF-sustained alpha RGC (A_2_), OFF-transient alpha RGC (A_3_), and pDSGC (A_4_) upon the addition of 20 µM histamine. (**B**) Background firing rates (averaged over 2 seconds before the appearance of the spot) before and after the application of histamine for the different RGC subtypes. Bold line indicates the mean±SEM, gray lines indicate paired cells. *p*<0.0001 for (B_3_), paired t-test; *p*=0.0231 for (B_4_), independent-sample t-test. (**C**) PSTHs of example RGCs showing their light responses to different sized spots (100, 400 and 800 µm) before (top, black) and after (bottom, orange) the histamine application. (**D-E**) Population data of the maximal firing rates (D) and response duration (log scale) (E) to different sized spots (mean±SEM). (D_3_) 200 µm, *p*=0.0389; 400, 600, and 800 µm, *p*<0.0001. (E_3_) 100 and 200 µm, *p*=0.0003 and 0.0346, respectively. (**E_4_**) 50 and 100 µm, *p*=0.0003 and *p*=0.0314, respectively. All two-way ANOVA with Bonferroni post-hoc correction. *: *p*<0.05, ***: *p*<0.001, ****: *p*<0.0001.

### Histamine broadens the directional tuning of DSGCs

The finding that histamine changes pDSGCs’ responses to stationary spot stimuli led us to investigate whether it also alters their responses to moving stimuli. We presented pDSGCs with moving gratings and bars while recording in cell-attached mode. In contrast to pDSGCs’ poor light responses to spots, they exhibited robust light responses to moving stimuli and maintained their preferred posterior direction after histamine application. However, in the presence of histamine, their directional tuning significantly broadened, as indicated by the normalized vector sum (NVS), which measures a cell’s tuning sharpness (see *Methods,* Figures S6A-D).

This effect of histamine was not specific to pDSGCs, as Ca^2+^ imaging data and MEA recordings showed that other DSGC subtypes similarly broaden their tuning to moving grating stimuli. Of the 32 RGCs (out of 247) classified as DSGCs in the Ca^2+^ imaging experiments, 13 broadened their directional tuning (41%) following the administration of histamine (20 µM). This percentage is significantly higher than in the control data set, where only 1 of the 19 DSGCs broadened its tuning (Figures S6E-H). Of the identified DSGCs in the MEA data set (43/485 RGCs), 41.9% (18/43) showed reversible changes in their responses to moving gratings (i.e., increased or decreased their evoked response and displayed an effective washout; see *Methods*). Histamine significantly and reversibly broadened the directional tunings of these cells (mean±SD NVS: 0.30±0.13, 0.21±0.15, 0.30±0.19 in pre, histamine and washout, respectively; Figures 5A and 5B), with 94.4% of them (17/18 cells, or 39.5% of all DSGCs) reducing their NVS. This is significantly more than in the control data set, where only 2.6% (1/39) of DSGCs reversibly changed their light responses (Figures 5A and 5B). Histamine reduced direction selectivity of DSGCs of all preferred directions, suggesting that it affects multiple DSGC subtypes (Figure 5C). We then investigated how histamine affects firing rates to moving gratings and found that histamine reversibly increased both the background firing rate of DSGCs (Figure S8E) and the evoked responses in all directions, including both the preferred and null directions (Figure 5D).

**Figure 5.**
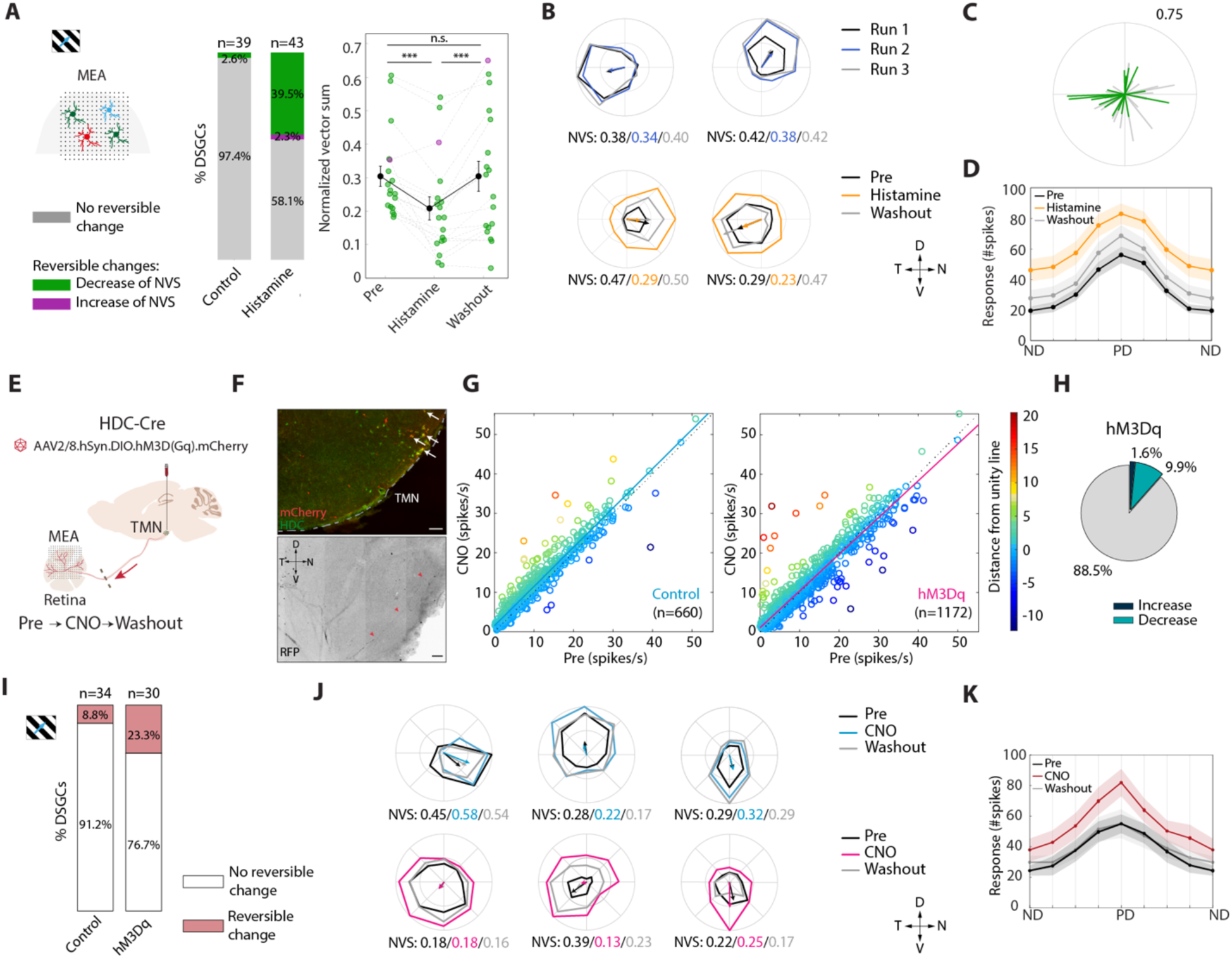
Histamine application, as well as pharmacogenetic activation of the histaminergic retinopetal axons, modulate DSGCs light-evoked response to moving gratings. (**A**) Responses of DSGCs to moving gratings (400 µm, 2 Hz) recorded on the MEA. Left: Percentages of non-reversibly changing cells (gray, 97.4% and 58.1% in control and histamine conditions, respectively, *p*<0.0001, Fisher’s exact test) and reversibly changing DSGCs, separated into cells that decreased (green) and increased (magenta) their NVS. Right: NVS of all the DSGCs that showed reversible changes in their light-evoked responses (n=18 cells) with mean±SEM overlaid (black; *p*=0.0003 (pre-histamine), *p*=0.0005 (histamine-washout), *n.s.* (pre-washout), Wilcoxon signed-rank test). (**B**) Top: Polar plots of example DSGCs classified as stable/not reversibly changing their evoked firing rate under control condition. Bottom: Example DSGCs that reversibly changed their evoked firing rate upon histamine application, accompanied with a decrease in directional tuning. Responses are normalized to the maximum response over three conditions. Arrows point to the preferred direction and their length represents the NVS. (**C**) Polar plot showing the preferred directions for all DSGCs that decreased their tuning with histamine (n=17 cells, green cells from (A)). The remaining DSGCs are shown in gray (n=26 cells). Arrow length represents the NVS. Retinal coordinates as in (B). (**D**) Mean±SEM response in all directions, aligned to the preferred direction, for all three conditions. Cells shown in (D) are green cells from (A), left. (**E**) Schematic representation of hM3Dq-mCherry microinjection into the TMN region of HDC-Cre transgenic mice. (**F**) Top: Immunostaining of mCherry and HDC in the TMN of injected mice, showing hM3Dq-mCherry^+^ cell bodies (red) and HDC^+^ neurons (green). Arrows point to examples of double-labeled neurons. Bottom: hM3Dq-mCherry^+^ histaminergic retinopetal axon (black, depicted by red arrowheads) in the wholemount dorsal retina, immunostained with an mCherry antibody (RFP) after MEA recording. Scale bars, 100 µm. (**G**) Firing rate before (x-axis) and after (y-axis) CNO drug application in control (left, non-injected mice) and hM3Dq (right) data sets. Individual RGCs are color-coded according to their distance from the unity line (*p*≪0.0001 and *p*≪0.0001; two-sided sign test). (**H**) Percentage of RGCs that showed an increase, decrease, or no significant change in their firing rate upon CNO application in hM3Dq data set (95% CI: 9.82% to 13.5%, *p*≪0.0001, binomial test). (**I**) Percentages of non-reversibly (white) and reversibly (red) changing DSGCs in control and hM3Dq, respectively. (**J**) As in (B) for control (top) and hM3Dq (bottom) experiments. (**K**) Mean±SEM response in all directions, aligned to the PD, for all conditions (n=7 cells). Abbreviations: D – dorsal, N – nasal, V – ventral, T – temporal. *: p<0.05, **: p<0.01, ***: p<0.001.

To characterize the source of the changes evoked in DSGCs, we targeted pDSGCs for intracellular recordings. Current-clamp recordings of the spiking activity showed that the histamine-induced increase in spontaneous firing rates (Figures S7A and S7B) was accompanied by a depolarized baseline voltage, which probably contributes to the enhanced firing (Figure S7E). In addition, histamine application broadened the spike shape and caused a reduced afterhyperpolarization (Figures S7C and S7D). This may imply that histamine changes the intrinsic properties of pDSGCs but does not preclude the option that it also acts on upstream neurons that, in turn, change the inputs to pDSGCs.

### Activation of histaminergic retinopetal axons in the retina modulates RGCs’ output

We next tested the contribution of histaminergic retinopetal axons to retinal processing by directly activating them chemogenetically (using a Designer Receptors Exclusively Activated by Designer Drug (DREADD)-based strategy) while recording RGC activity. We injected a AAV2/8.hSyn.DIO.hM3D(Gq)-mCherry, which drives the expression of the hM3Dq receptor in a Cre-dependent manner, into the TMN of HDC-Cre mice (hereafter referred to as hM3Dq (Gq-coupled human M3 muscarinic DREADD)). Between 4-6 weeks after the injection (Figures 5E and 5F), we carried out MEA recordings from isolated retinas of hM3Dq-injected HDC-Cre mice to assess changes in RGC activity following clozapine-N-oxide (CNO) bath application, which activates the hM3Dq receptor (Alexander et al., 2009; Armbruster et al., 2007; Rogan and Roth, 2011; Roth et al., 2016). As a control, we performed the same experiments on non-injected mice. We first recorded the baseline and light-evoked activity to a set of visual stimuli under perfusion with control solution (pre) for ∼30 minutes before adding 15 µM CNO, and then repeated the same visual stimuli protocol (∼30 min), followed by washing out the CNO and repeating the same set of visual stimuli a third time (∼30 min). The CNO washout and a third repeat were added to investigate whether the effects caused by CNO are transient or can persist after the drug is removed.

Our findings show that CNO administration induces a change in the baseline firing rate of a portion of RGCs in both the control and hM3Dq experimental groups (Figure 5G and Figure S8C; hM3Dq and control, n=1172 (13 retinas) and 660 (5 retinas) RGCs). As the firing-rate distributions between the two data sets prior to adding CNO were not significantly different (Figure S8A), we calculated the averaged firing rate difference from the respective pre-condition (Δ firing rate, CNO-pre) and compared the cumulative distribution functions. The distributions of the change in firing rate differed significantly between the hM3Dq and control data sets (Figure S8B). Overall, a significantly greater proportion of RGCs from hM3Dq retinas were responsive to CNO application. Of the 1172 RGCs recorded, 11.5% (*n*=135/1172) showed either increased or decreased spiking activity (in comparison, only 4.8% (n=32/660) of RGCs increased or decreased their spiking activity in the control, Figure 5H). Out of the RGCs that reacted to CNO application in hM3Dq data sets, 85.9% (*n*=116/135) reduced their firing rates and only 14.1% (*n*=19/135) increased it upon the introduction of CNO (see *Discussion*).

Next, we sought to determine the effects of chemogenetic activation of histaminergic retinopetal axons on light-evoked RGC activity. Given the results we obtained regarding DSGCs in the bath application of histamine, we focused on this population. We identified DSGCs based on their responses to moving gratings and obtained 34/698 and 30/1043 RGCs from 5 and 11 retinas in control and hM3Dq experiments, respectively. As before, we included only RGCs with reversible changes in their evoked spiking activity to moving gratings (see *Methods*). While only 3/34 (8.8%) DSGCs showed reversible changes in control, 7/30 (23.3%) showed reversible changes in hM3Dq experiments (Figure 5I). The DSGCs that reversibly changed their light-evoked response in hM3Dq either increased or decreased their NVS (Figure S8D and Figure 5J), as opposed to the histamine bath application results (Figure 5A). Yet, the DSGCs in the hM3Dq experiments increased their firing rates in all directions, including the ND and PD (Figure 5K), similarly to their behavior in the histamine application experiments (Figure 5D). An increase in spiking activity was also observed in their firing rates recorded before presenting the moving grating stimulus (Figure S8F).

### Histamine enhances DSGCs’ responses to high motion velocities

Running speed is a common measure for arousal (McGinley et al., 2015). Given that histaminergic neurons are known to increase their firing rate with arousal, we considered the possibility that histamine shifts the velocity tuning of DSGCs to favor higher velocities, resulting from increased running speed, at the expense of tuning sharpness (Figures 5 and S6). To test this, we patch-clamped pDSGCs, and presented them with moving gratings at various temporal frequencies, ranging from 1 to 12 Hz (corresponding to 400 and 4800 µm/s), instead of only 2 Hz as before (Figures 5 and S6). We assessed their responses before and after histamine application (10 µM). In control conditions, pDSGCs were tuned to lower speeds, exhibiting high discharge rates in response to slow-moving gratings (1-2 Hz) and diminished or no responses to faster moving gratings (4-12 Hz; Figures 6A-C). Upon histamine application, pDSGCs improved their ability to encode faster-moving stimuli as is evident by the significant increase in the normalized response in the preferred direction at 6 and 8 Hz (Figures 6A-C), while maintaining direction selectivity at these high motion velocities (Figure 6D). This increase in firing rate was not merely a byproduct of the increased spontaneous activity, since the responses were time-locked to the grating stimulus (Figure 6B). To further investigate this, we examined the first harmonic magnitude (extracted at the stimulus frequency) computed by fast Fourier transform (FFT) of the mean PSTH in response to the PD motion in a larger dataset that contained different subtypes of DSGCs recorded on the MEA (n=27). We observed a significant increase in the FFT magnitude for responses to gratings moving at 4, 6 and 8 Hz after histamine application (Figures 6E and 6F). Together, these findings show that histamine extends the range of motion speeds that DSGCs can encode, enabling them to track faster moving stimuli in a time-locked manner.

**Figure 6.**
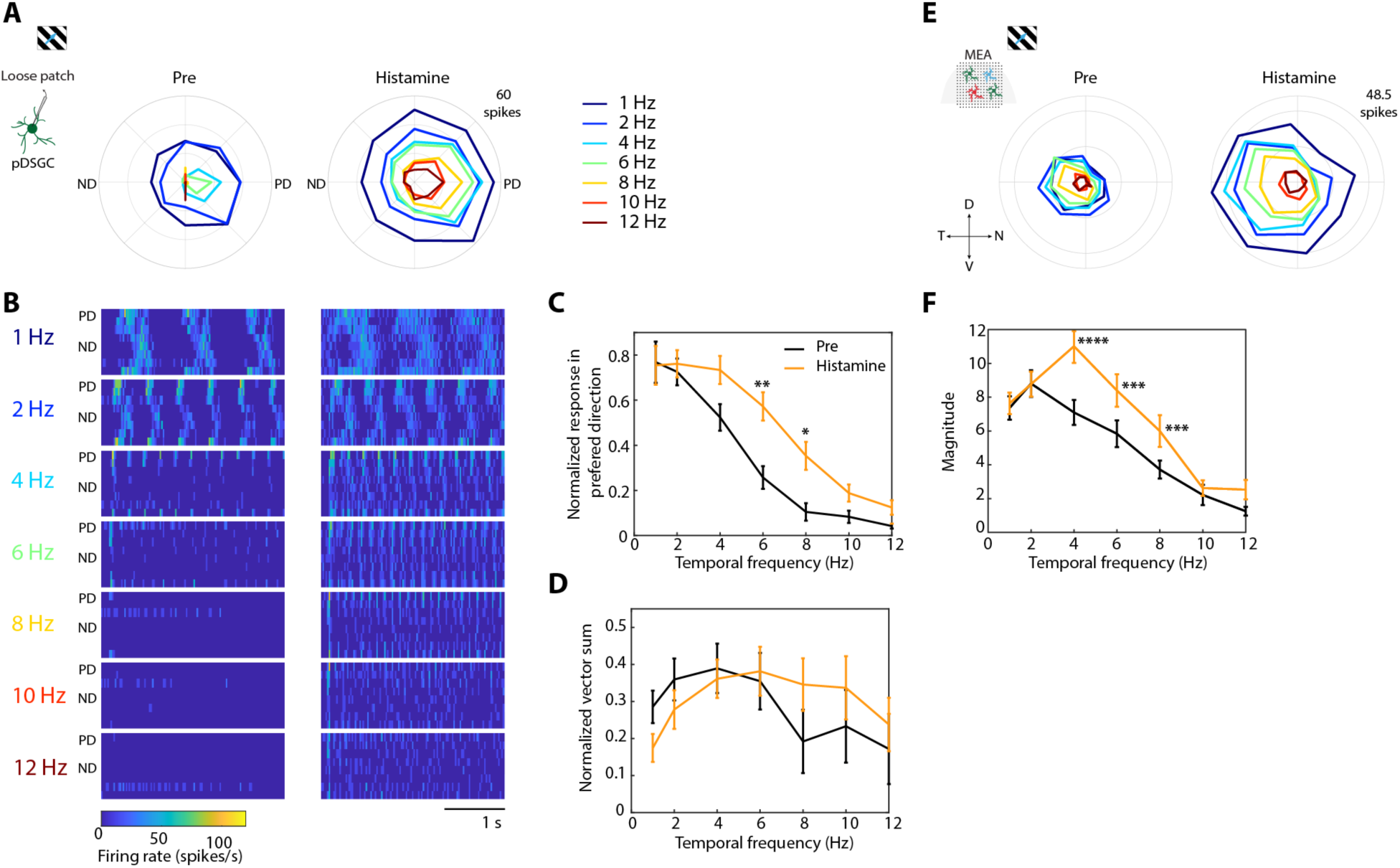
Histamine enhances DSGCs’ responses when presented with higher velocity stimuli. (**A**) Polar plots showing the responses of an example pDSGC to moving gratings at different temporal frequencies, obtained using patch clamp recordings, before (left) and after 10 µM histamine application (right). PD, preferred direction; ND, null direction. Colors denote temporal frequencies. (**B**) Heatmaps showing the mean firing rates of the cell in (A) in response to different temporal frequencies, before (left) and after (right) histamine application. Rows correspond to the eight different directions of the grating stimulus (PD and ND according to (A)). (**C-D**) Population data (mean±SEM) of the normalized response in the preferred direction (C, *p*=0.0013 for 6 Hz and *p*=0.025 for 8 Hz) and normalized vector sum (D). n=9-11 cells for each frequency. Plots for before (pre) and after histamine are shown in black and orange, respectively. (**E**) Same as (A) for a representative DSGC recorded on the MEA. (**F)** First harmonic magnitude of the FFT at the stimulus frequency as a function of temporal frequency for all DSGCs recorded on the MEA (n=27; *p*<0.00001, *p*=0.0001 and *p*=0.00097 for 4, 6 and 8 Hz, respectively). Two-way ANOVA with Bonferroni post-hoc correction. Abbreviations: D – dorsal, N – nasal, V – ventral, T – temporal. *: *p*<0.05, **: *p*<0.01, ***: *p*<0.01, ****: *p*<0.0001.

### H_1_R antagonist non-uniformly affects human’s sensitivity across the visual field

Having established a role for histaminergic retinopetal axons in mice, we wondered if histamine may also affect visual information processing in the human retina, given that retinopetal axons have been found in human retinas (Honrubia and Elliott, 1968; Repérant et al., 1989), and histaminergic retinopetal axons in non-human primates (Gastinger et al., 1999). To study the effect of histamine on human visual sensitivity, we administered the first-generation H_1_R antagonist dimetindene maleate (Fenistil) to 8 human volunteers. The participants underwent a series of visual tests twice, once after taking the H_1_R antagonist and once after taking a placebo, in a single-blind experimental design (see *Methods*). The H_1_R antagonist did not affect the subjects’ levels of concentration, as assessed by total time of the visual tests (Table S1).

We first performed a visual field test of the central visual field. Subjects were shown spots of light at different intensities at 68 different locations within 10° of the fovea. A visual threshold was then calculated in decibels (db) for each location (see *Methods*). The difference in visual sensitivity between the H_1_R antagonist and placebo revealed a slight decrease in nearly all locations (Figure 7a). Averaging over all locations for each eye revealed a slight but significant decrease in sensitivity with the H_1_R antagonist (Figure 7B). Next, we examined the visual sensitivity in the peripheral visual field (between 30°-60°; Figure 7C). The difference in visual sensitivity between the H_1_R antagonist and placebo revealed an opposite trend in different locations in the visual field, with the H_1_R antagonist causing increased visual sensitivity in the superior field and decreased sensitivity in the inferior field (Figures 7C and 7E). Thus, we calculated the mean visual sensitivity in four locations in the visual field (superior, inferior, nasal and temporal), and found that the H_1_R antagonist significantly increased visual sensitivity in the superior region (Figure 7D). Our findings suggest that histamine may non-uniformly affect the human retina, decreasing the sensitivity of the ventral retina, which represents the superior visual field, while possibly increasing that of the dorsal and central retina, which represent the inferior and central visual field, respectively.

**Figure 7.**
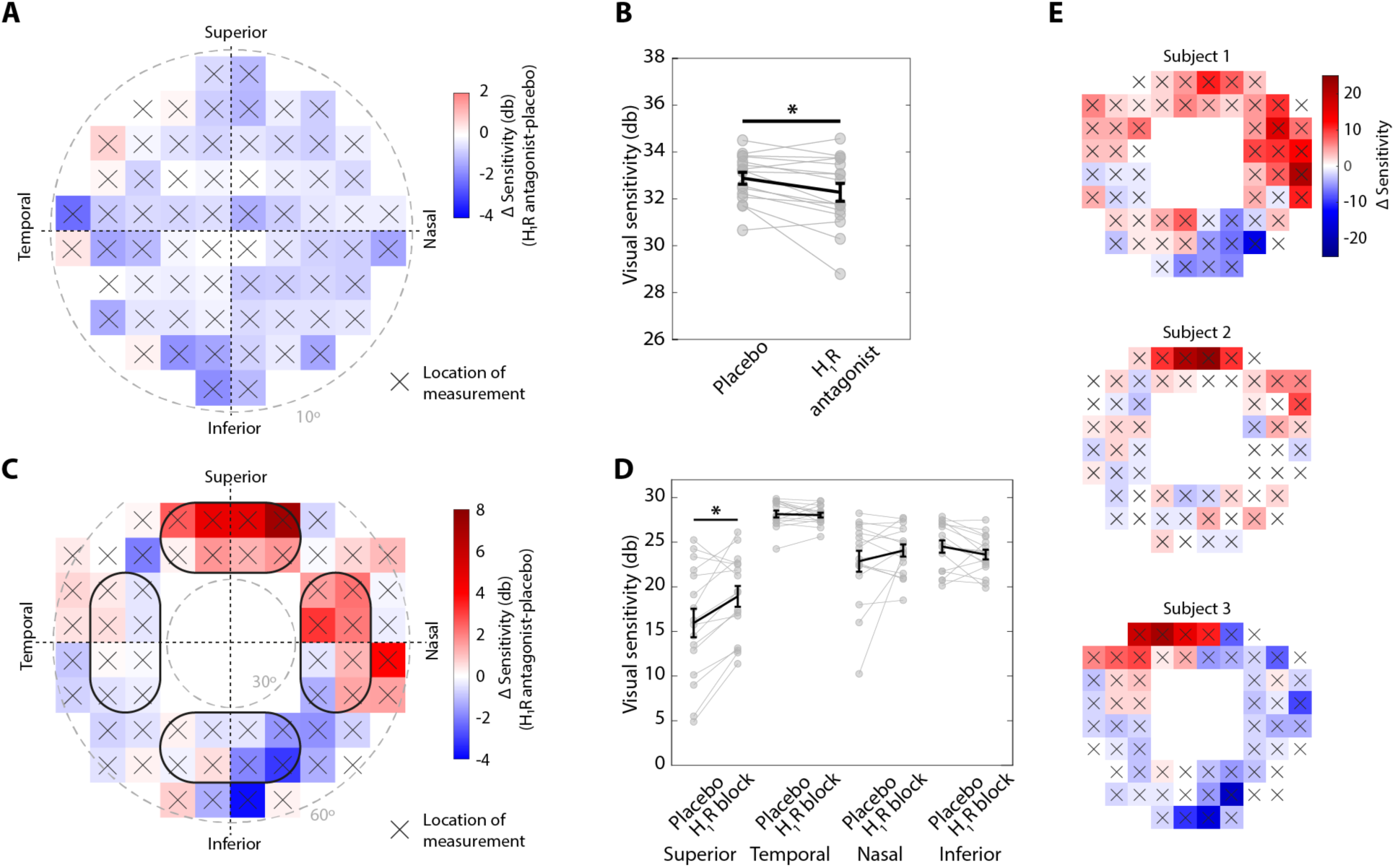
H_1_R antagonist non-uniformly affects visual sensitivity in the peripheral visual field in humans. (**A**) Heatmap showing the mean change in decibels (db) at each measured location in the central visual field test. The center corresponds to the fovea. Positive (red) and negative (blue) values correspond to increased and decreased sensitivity, respectively, in the presence of the H_1_R antagonist compared to the placebo. Gray, dashed line labels 10° from the fovea. (**B**) Visual threshold sensitivity with placebo and with H_1_R antagonist, calculated by averaging across all locations in the central visual field test (*p*=0.0150, two-tailed paired t-test). (**C**) As in (A) for the peripheral visual field test. Gray dashed lines label 30° and 60° of the visual field. (**D**) Visual threshold sensitivity with placebo and H_1_R antagonist for each region of the visual field, calculated by averaging across 8 locations (outlined in (C)). *p*=0.0102, two-tailed paired t-test with Bonferroni-Holm correction. (**E**) Like (C), but for three example subjects. *: *p*<0.05, n=16 eyes from 8 subjects.

## Discussion

In this study, we sought to determine how histaminergic retinopetal projections shape retinal processing. We demonstrated that histaminergic projections originating in the TMN of the hypothalamus innervate the retina and can shape the response properties of distinct RGC subtypes. Using both two-photon Ca^2+^ imaging and MEA recordings, we showed that histamine increases the baseline activity and qualitatively changes the light responses of many RGCs. We focused on specific RGC subtypes, and using patch-clamp recordings, we showed that while the activity in ON-sustained and OFF-sustained alpha RGCs remains unaltered, histamine shortens the light responses of OFF-transient alpha RGCs and improves pDSGCs’ ability to respond to higher velocity stimuli, while broadening their directional tuning at lower velocities. This broadening of directional tuning was also exhibited by other DSGC types that were recorded using Ca^2+^ imaging and MEA recordings. Furthermore, direct activation of histaminergic projections using chemogenetic techniques elicited changes in the spontaneous and light-evoked spiking activity of RGCs, including DSGCs. Finally, we found that administration of an antihistamine non-uniformly affects humans’ visual sensitivity, suggesting that this pathway is evolutionarily conserved across species. Our results reveal a top-down functional circuit of brain-derived histaminergic projections that shape visual processing in the earliest possible site – the retina.

### Modulation of visual processing by arousal state

The brain’s histaminergic system has long been associated with wakefulness and attention (Haas et al., 2008; Monnier et al., 1970; Scammell et al., 2019; Wada et al., 1991). Histaminergic neurons are silent during sleep, and their firing rate increases with the arousal state of the animal, peaking during attentive waking (Takahashi et al., 2006). As our results demonstrate that histamine modulates both the basal firing and the light responses of RGCs, we can hypothesize that visual processing can change with the arousal state. Indeed, neurons in V1, dLGN and SC were shown to change their spontaneous and visually driven activity with the level of arousal (Aydın et al., 2018; Erisken et al., 2014; Ito et al., 2017; Mineault et al., 2016; Niell and Stryker, 2010; Roth et al., 2016; Savier et al., 2019; Stringer et al., 2019; Vinck et al., 2015; Williamson et al., 2015). The origin of these changes, however, is usually attributed to local neuromodulators, top-down circuits, or local connectivity. Notably, two recent studies revealed that the retinal output itself changes with the mouse’s arousal state both in the dLGN (Liang et al., 2020) and in the SC (Schröder et al., 2020). Using *in vivo* Ca^2+^ imaging of RGCs’ axonal boutons, these studies demonstrated that visual responses of almost 50% of them are modulated with locomotion and pupil size, which reflect the arousal state. Typically, these modulations included suppression of visual responses and attenuation of direction and orientation selectivity, although this was only shown for low motion velocity (Liang et al., 2020; Schröder et al., 2020). Interestingly, neurons in the primary auditory cortex also broaden their frequency tuning as arousal increases (Lin et al., 2019).

The arousal-related suppression of visual responses and direction selectivity reported in both the dLGN and SC (Liang et al., 2020; Schröder et al., 2020) is in line with our findings that histamine shortens the responses of OFF-transient alpha RGCs and broadens the tuning of DSGCs in response to low velocity motion. Yet, several aspects of the arousal modulation differ between the dLGN and SC. These differences could originate from different subtypes of RGCs that innervate the dLGN and SC (Ellis et al., 2016), but they may also have other origins. Indeed, other studies have shown that local presynaptic modulation also influences the effects of arousal on the activity of retinal axonal boutons (Chen and Regehr, 2003; Lam and Sherman, 2013; Seeburg et al., 2004; Yang et al., 2014). Since RGCs project onto numerous brain targets to transfer the visual information (Martersteck et al., 2017; Morin and Studholme, 2014), we hypothesize that properties of arousal modulation that contribute to all the targets would take place at the retina, while modulations that serve a specific target would take place locally.

### Species-specific effects of histamine on retinal cells

Several studies investigated the effects of histamine application on retinal neurons’ activity *ex vivo*. Histamine was found to act on cones and bipolar cells in primates (Vila et al., 2012; Yu et al., 2009) and to enhance the activity of various amacrine cells in mice (Gastinger et al., 2006b; Horio et al., 2018; Yu et al., 2009). According to two reports, histamine affects >80% of the RGCs, but in a highly variable manner (Akimov et al., 2010; Gastinger et al., 2004). In primates, histamine either increases or decreases the spontaneous activity of RGCs, while typically suppressing their light-evoked responses (Akimov et al., 2010; Gastinger et al., 2004). In rats, histamine increases the spontaneous activity of most affected RGCs, with variable effects on their light-evoked responses (Gastinger et al., 2004), similar to our findings in the mouse retina. Taken together, these studies suggest that histamine’s effects may vary between species. In both primates and rats, different histamine concentrations (1-50 µM) caused similar trends, but effects were stronger with the higher concentration tested (Gastinger et al., 2004). Our results similarly show a histamine dose-dependent effect on RGCs activity.

### Mechanism of histamine-induced modulation

While immunohistochemical studies have revealed the presence of H_1_R in rat and mice dopaminergic amacrine cells, this receptor is not restricted to these cells, as it is expressed also in all layers of the inner plexiform layer and in the ganglion cell layer (Gastinger et al., 2006b; Greferath et al., 2009). H_2_R expression was observed in Müller glia cells and H_3_R could not be localized, which may have been due to technical issues (Greferath et al., 2009). In our search for the expression levels of histamine receptors in recent single-cell RNA sequencing data sets of the mouse retina (at P14-19 (Macosko et al., 2015; Shekhar et al., 2016; Yan et al., 2020) and adulthood (Tran et al., 2019)), using the Single Cell Portal (SCP, Broad Institute), we noticed that H_3_R mRNA transcripts were detected in all major neuronal subtypes of the retina (Macosko et al., 2015), H_2_R mRNA transcripts only in RGCs and a subpopulation of amacrine cells, and H_1_R mRNA transcripts in RGCs and subsets of bipolar and amacrine cells (Macosko et al., 2015; Shekhar et al., 2016; Tran et al., 2019; Yan et al., 2020). Our results demonstrate that histamine-induced increases in RGCs’ baseline activity are primarily mediated by H_1_R (Figure 2G). Yet, it is possible that H_2_R and H_3_R are also involved in histamine-induced changes in RGCs’ light responses. Regardless of the receptors involved, the effects of histamine are probably the result of a combination of actions on various interneurons that modulate the inputs to RGCs as well as intrinsic RGC properties, as suggested by our intracellular recordings, which revealed a change in the spike width of pDSGCs following the histamine application (Figure S7).

### Behavioral consequences of histamine-induced retinal changes

In all our AAV tracing experiments in which we tracked the retina’s orientation, the retinopetal axons projected primarily to the dorsal retina, despite using several different AAV serotypes and promoters and two HDC-Cre mouse lines (see *Methods*). These retinopetal axons arise from just a few neurons (2-3 neurons, determined by the number of axons labeled in the optic nerve), so although it is possible that we failed to transfect all histaminergic retina-projecting neurons, it is reasonable to conclude that their number is low, and that they favor the dorsal retina.

Indeed, it is known that the mouse retina is not uniform, as the density of UV cones is higher in the ventral retina and the density, morphology and even physiology of distinct RGC subtypes differs between the dorsal and ventral retinal halves (reviewed in Heukamp et al. (2020)). These topographic variations may result from the different functional requirements of the two retinal halves. In the case of the mouse, a small animal that views the environment from close to the ground, its upper and lower visual fields primarily correspond to the sky and the ground, falling on the ventral and the dorsal retina, respectively. As a result, the two retinal halves process different visual statistics (Qiu et al., 2021) and exhibit different functional specializations: while the dorsal retina is involved in foraging, the ventral retina is mainly concerned with predator detection and even hunting (Heukamp et al., 2020; Johnson et al., 2021). As the mouse begins to move through its environment, the optic flow will increase non-uniformly across the visual field, with higher apparent velocities for objects in the lower (and closer) visual field, encoded by the dorsal retina. As the histaminergic retinopetal axons innervate the dorsal retina and the release of histamine is modulated by the arousal state, it could be hypothesized that these axons fine-tune retinal processing during periods of high arousal, shifting the velocity tuning of DSGCs to encode faster motion, which may be beneficial during running (e.g. escaping or foraging). Since the ventral retina lacks histaminergic innervation, the responses of ventral RGCs are expected to be unaltered *in vivo*, keeping the ability of the mouse to detect predators in the upper visual field.

### Histaminergic retinopetal axons modulate RGC activity

When we directly activated histaminergic retinopetal axons *ex vivo*, using chemogenetics, a greater percentage of RGCs from injected mice altered their background activity following CNO application compared with the control group. Unlike the effects of histamine bath application, which tended to increase the firing rate of RGCs, the chemogenetic activation of histaminergic fibers caused, on average, a decrease in the firing rate of RGCs, although a small portion of the RGCs did significantly increase their activity. The explanation for this discrepancy may come from the fact that a portion of histaminergic neurons co-transmit histamine and GABA in a paracrine manner (Kukko-Lukjanov and Panula, 2003; Yu et al., 2015). It remains unknown whether histaminergic retinopetal axons also exhibit this property, but if this is indeed the case, stimulating retinopetal axons would result in a more complex circuitry activation, potentially suggesting a synergetic role for GABA and histamine in shaping the retinal code according to specific behavioral requirements. Additionally, we cannot exclude the possibility that our findings result from a combination of hM3Dq receptor activation on histaminergic axons and potential off-target effects of CNO (Gomez et al., 2017; Jendryka et al., 2019), which were indeed observed in the control group. Particularly, as one of CNO’s off-target effects is the inhibition of H_1_R binding, it is possible that the changes in RGCs activity we detected in the chemogenetic experiments were primarily mediated via H_2_R and H_3_R, a factor that, too, could contribute to the differences from histamine bath application experiments.

Nevertheless, similarly to the results obtained following the pharmacological histamine application, we found a subpopulation of DSGCs that increase their firing rate in response to moving gratings without changing their directional preference, gradually returning to pre-stimulation levels after the washout. These findings suggest that at least part of our observations with the histamine bath application can be replicated using chemogenetic activation of histaminergic fibers.

### Histaminergic modulation of visual sensitivity in humans

To investigate how an H_1_R antagonist may affect human visual sensitivity, we administered dimetindene maleate (Fenistil) orally, which allowed Fenistil to potentially reach not only the retina, but also other brain structures innervated by histaminergic axons. This makes it challenging to disentangle the H_1_R antagonist’ effects on the retina from those on the brain, including drowsiness (Mahdy and Webster, 2017). However, several lines of evidence suggest that the effects we observed on visual sensitivity are indeed the result of the H_1_R antagonist acting at the level of the retina. First, test time was not increased by the H_1_R antagonist, confirming that concentration was not affected by drowsiness. Second, our findings revealed a non-uniform effect in different locations of the visual field: decreased sensitivity in the center and inferior areas of the visual field and increased sensitivity in the superior field. We postulate that if the actions of the H_1_R antagonist had occurred downstream to the retina, we may have seen a more uniform effect across the visual field. Notably, the flash sensitivity of baboon RGCs recorded *ex vivo* was shown to decrease with histamine application (Akimov et al., 2010), in line with our findings.

While it is not possible to administer histamine to human subjects, we hypothesize that this would cause opposing effects, namely, an increase in visual sensitivity in the central and inferior visual field and a decrease in sensitivity in the superior visual field. As the superior visual field has the lowest visual sensitivity to begin with (Figure 7D), this suggests that humans rely less on the superior visual field and that, like in mice, histamine can tune visual processing to selectively enhance sensitivity in specific areas according to behavioral needs. H_1_R antagonists were previously shown to decrease the critical flicker fusion frequency in humans (Nicholson and Stone, 1983; Nicholson et al., 1982; Turner, 1968), suggesting that histamine also affects temporal sensitivities in the retina. Since these studies, as well as ours, only tested a selective H_1_R antagonist, while all three histamine receptors are expressed in the primate retina (Gastinger et al., 2006b; Vila et al., 2012), histamine’s effects on retinal processing may be even more complex.

### Neuromodulators in the visual system

Over the years, the search for mammalian retinopetal axons, and particularly their origin, has led to opposing findings, even within species. Studies based on axonal tracers reported various origins, including the hypothalamus, various visual structures, the oculomotor nucleus and the dorsal raphe nucleus (Bons and Petter, 1986; Hoogland et al., 1985; Itaya and Itaya, 1985; Labandeira-Garcia et al., 1990; Larsen and Møller, 1985; Terubayashi et al., 1983; Villar et al., 1987). Other studies failed to label any brain area or interpreted somatic labeling in the brain as the result of transneuronal transport (Davis and McKinnon, 1982; Repérant, 1975; Schnyder and Künzle, 1984; Weidner et al., 1983)(Davis and McKinnon, 1982; Repérant, 1975; Schnyder and Künzle, 1984; Weidner et al., 1983). In most of these investigations, the axonal tracers labeled only a few cell bodies, which contributed to the difficulty of finding the retinopetal axons and their origin. Here, we took advantage of the HDC-Cre mouse lines to indisputably identify histaminergic neurons in the TMN as a source for retinopetal axons. Future work may make use of other transgenic mouse lines to resolve whether other brain regions also contribute to visual processing in the retina. Particularly, it was suggested that serotonergic neurons in the dorsal raphe nucleus send projections to the retina (Gastinger et al., 2005; Repérant et al., 2000; Schütte, 1995). If true, this suggests that the histaminergic and serotonergic systems, which contribute to higher cognitive functions, including wakefulness and mood, may interact already at the level of a primary sensory organ.

## Acknowledgements

We thank Hillary Voet for statistical consultation, Maya Groysman for custom viral production, Mark Shein-Idelson for sharing the framework for the visual stimulation GUI, Meital Oren-Suissa for using her confocal microscope, Efrat Solomon for help with histology, and members of the Gollisch lab for helpful advice about MEA recordings. This work was supported by research grants from the European Research Council (ERC starter No. 757732), the Israel Science Foundation (2449/20), the Minerva Foundation with funding from the Federal German Ministry for Education and Research, and by the Charles and David Wolfson Charitable Trust, Rolf Wiklund and Alice Wiklund Parkinson’s Disease Research Fund, Consolidated Anti-Aging Foundation, and Dr. Daniel C. Andreae. A.H. was supported by the Minerva Fellowship, L.A. is supported by ISEF, S.R. is supported by a Dean Fellowship of the Weizmann Institute of Science, M.R.-E. is incumbent of the Sara Lee Schupf Family Chair.

## Methods

### Mice models and ethical statement

Two-photon targeted recordings from DSGCs were performed using Trhr-EGFP mice (MMRRC, stock #030036-UCD), which express GFP in posterior-preferring ON-OFF DSGCs (Rivlin-Etzion et al., 2011). Two-photon Ca^2+^ imaging and two-photon targeted recordings from alpha-RGCs were conducted from the ganglion cell layer of the isolated retina of mice expressing GCaMP6f in RGCs (Jackson laboratory, stock #025393 (Dana et al., 2014)(Dana et al., 2014)). Intracranial injections were performed on C57Bl/6J mice (purchased from Charles River Breeding Laboratories) and on two HDC-Cre mouse lines, which express Cre recombinase under the control of the *hdc* promoter (Jackson laboratory, stock #021198 (Zecharia et al., 2012) and MMRRC, stock #037409). MEA experiments were performed on wildtype mice from the same colony. Weaned mice from either sex, 4-12 weeks old, were housed in groups of no more than five in individual cages at 25°C in a 12h/12h light-dark cycle with water and food provided *ad libitum*. All procedures were approved by the Institutional Animal Care and Use Committee (IACUC) at the Weizmann Institute of Science.

### Intracranial AAV injections

To label retinopetal axons in C57Bl/6J mice, 0.5 to 1 µl of AAV2/8.hSyn.mCherry (HUJI Vector Core #7.19) or AAV2/8.hSyn.Chronos.tdTomato (Addgene #62726) were injected into the TMN based on stereotactic coordinates (injection site: antero-posterior (AP) = -2.6 mm; medio-lateral (ML) = 0.7 mm; dorso-ventral (DV) = -5.3 mm from bregma). To label histaminergic retinopetal axons in HDC-Cre mice, we used Cre-dependent adeno-associated viruses (AAV2/1.CAG.Flex.tdTomato.WPRE.SV40 or AAV2/8.CAG.Flex.tdTomato.WPRE.SV40, Harvard Vector Core, Lot No. 704 and 605, respectively). For chemogenetic electrophysiological experiments, mice were unilaterally or bilaterally injected in the TMN with AAV2/8.hSyn.DIO.hM3D(Gq)-mCherry (HUJI Vector Core #35.18).

For the injection procedure, mice were anesthetized with inhalant isoflurane (5% induction and 1.5-2% maintenance, SomnoSuite, Kent Scientific) and administered 0.5 ml saline via intraperitoneal injection, to avoid dehydration. The animal was kept on a closed loop heating pad and watched for vitals throughout the surgery and its eyes were kept from drying with a layer of Synthomycine (ABIC Ltd., TEVA Pharmaceutical Industries Ltd., Israel). Next, a craniotomy of 1-2 mm was made, 2.5-2.8 mm posterior and 0.7-1 mm mediolateral to Bregma. A Hamilton syringe (1 µl, 65458-01) was then lowered into the brain at a rate of 10 µm/s to target the TMN. The viral solution was delivered at 0.1 µl/min after a 10 min pause to allow the brain to resettle. The scalp incision was sealed with a tissue adhesive (Histoacryl, Melsungen AG, Germany) and mice were left to recover post-surgery, after two subcutaneous injections of antiseptic analgesia (0.01xNorocarp, Norbrook Laboratories Limited, Newry, Co. Down, Northern Ireland, 10 µl/gr).

### Tissue processing, immunohistochemistry protocols and microscopy

Immunohistochemical analysis of virus expression (reporter gene, mCherry or tdTomato) in combination with the identification of histaminergic neurons via HDC immunohistochemistry was performed on mouse brain slices 3-4 weeks after the injection (primary antibody: rabbit polyclonal anti-histidine decarboxylase, 1:300, PROGEN Biotechnik GmbH, Cat. No. 16045; secondary antibody: donkey anti-rabbit 488, 1:200, Invitrogen A21206). Mice were deeply anesthetized with a terminal intraperitoneal injection of pentobarbital (Pentobarbital Sodium, 200 mg/ml, CTS Chemical Industries Ltd., Kiryat Malachi, Israel), then intracardially perfused with phosphate buffered saline (PBS, Biological Industries Israel, 02-023-1A, pH 7.4) and 4% paraformaldehyde (PFA, ChemCruz, Santa Cruz Biotechnology, Inc., CAS: 30525-89-4) before brain and eye extraction. Eyecups were fixed for 1 hour (4% PFA) and then hemisected to obtain wholemount retinas.

Brains were fixed further for 24-48 h in 4% PFA and washed in PBS, then sliced (30 µm) by a vibratome (7000 smz-2 Vibratome, Campden Instruments Ltd.). Slices were washed three times in PBS and subsequently blocked with 0.25% PBST (PBS + Triton X-100, Sigma, CAS: 9002-93-1) with 3% bovine serum albumin (BSA, MP Biomedicals, Cat no. 160069) for 2 hours at room temperature, followed by overnight immersion in primary antibody solution (1% BSA and 0.1% Triton X-100 in PBS with antibody-specific dilution) at 4°C on a shaker. The next day, slices were washed in PBS and immersed in a secondary antibody solution overnight (1% BSA in PBS with antibody-specific dilution). Slices were mounted onto Superfrost/Plus Microscope Slides (Thermo Scientific), covered with a coverslip using Vectashield antifade mounting medium with DAPI (Vector laboratories, H1200). All brain sections were digitally scanned using Olympus UPlanSApo 10x/0.40 NA or 20x/0.75 NA objectives on an Olympus BX61VS slide scanner (Olympus Corporation, Tokyo, Japan).

Retinal wholemounts were blocked with 0.25-0.4% PBST with 3-5% BSA for 2 hours at room temperature and then incubated for two days in primary antibody solution (1% BSA and 0.1% Triton X-100 in PBS; primary antibodies: goat polyclonal anti-RFP antibody, 1:300, MyBioSource, Cat. No. M5448122 or rabbit polyclonal anti-histamine 1 receptor, 1:100, Alomone labs, Cat. No. #AHR-001) at 4°C on a shaker. The next day, slices were washed in PBS and overnight immersed in a secondary antibody solution (1% BSA in PBS with antibody specific dilution; secondary antibodies: donkey anti-goat 568, 1:1000, Invitrogen, Cat. No. #A11057, or donkey anti-rabbit 546, 1:1000, Invitrogen, Cat. No. #A10040). The tissues were stained with DAPI to identify nuclei and mounted onto Superfrost/Plus Microscope Slides, covered with a coverslip, using a Vectashield antifade mounting medium with DAPI. Retinal wholemounts were imaged using an inverted laser scanning confocal microscope (Zeiss, Oberkochen, Germany) equipped with 488, 543, and 633 nm laser lines, using ZEN software (Zeiss). Z-stack images were acquired using a 63x/1.4 Plan Apochromat oil objective with a step size of 0.25 µm. Tiled images of the whole retina were acquired using a 20x/1.0 W Plan Apochromat DIC VIS-IR 75 mm objective. Further image processing for brain slices and wholemount retinas was performed with Fiji software (Schindelin et al., 2012).

### Tissue preparation for physiology

Mice were kept in dark-adapted conditions for at least 30 min and then anesthetized with isoflurane (Terrell, Piramal Critical Care Inc.) and decapitated. Eyes were immediately enucleated and dissected under dim red and infrared light in a Petri dish containing Ames medium (Sigma, St. Louis, MO, USA) supplemented with 1.9 g/l of sodium bicarbonate saturated with carboxygen (95% O_2_ and 5% CO_2_). The orientation of the retina was determined based on landmarks on the choroid as described previously (Wei et al., 2010), and retinas were dissected in two halves along the nasal-temporal axis. Retinas were kept in the dark at room temperature in Ames’ medium bubbled with 95% O_2_/5% CO_2_ until used.

For MEA recordings, MEAs were precoated with poly-D-Lysine solution (PDL, 1.0 mg/ml in H_2_O, Merck-Millipore, CAT: A-003-E) for one hour at room temperature. After washing off the PDL, one half of the retina (chemogenetic experiments were performed only on dorsal retinas) was mounted on the multi-electrode array with the RGC layer facing the electrodes, as previously described in (Karamanlis and Gollisch, 2021). For targeted patch-clamp recordings, retinas were cut into half, isolated from the pigment epithelium, and mounted, photoreceptor side down, over a hole of 1–1.5 mm^2^ on a filter paper (GSWP01300, Merck Millipore, Billerica, MA, USA). For two-photon Ca^2+^ imaging, retinal pieces were mounted onto poly-D-Lysine-coated 12 mm coverslips (Product Number 354086, Corning, Glendale, AZ, USA).

### Histamine application and pharmacology

Histamine (Sigma-Aldrich, product number H7250)-containing Ames solution was prepared fresh from powder for each experiment (5-20 µM). Histamine receptor blockers, Cetirizine dihydrochloride (H_1_R; Tocris, Bristol, UK, product number: 2577) and JNJ 5207852 (H_3_R; Tocris, product number: 4020) were dissolved in water to make stock solutions of 20 mM and then were further diluted in Ames solution to a working concentration of 20 µM. Histamine receptor blocker famotidine (H_2_R; Sigma-Aldrich, St. Louis, MO, USA, product number: F6889) was dissolved in DMSO to make a stock solution of 80 mM and then was further diluted in Ames solution to a working concentration of 40 µM.

### Targeted patch-clamp recordings

Retinas were placed under a two-photon microscope (Bruker, Billerica, MA, USA) equipped with a Mai-Tai laser (Spectra-physics, Santa Clara, CA USA) and perfused with oxygenated Ames medium at 32-34°C. Identification of and recording from GFP^+^ RGCs were carried out as previously described (Rivlin-Etzion et al., 2011; Wei et al., 2010). In short, GFP^+^ cells were identified using the two-photon microscope laser at 920 nm, to avoid bleaching of the photoreceptors. Alpha RGCs were targeted in GCaMP6f mouse retinas by finding RGCs whose cell bodies had a diameter greater than 20 µm (van Wyk et al., 2009); pDSGCs were targeted in Trhr-EGFP mice retinas. The inner limiting membrane above the targeted cell was dissected under the microscope with a glass electrode using infra-red illumination. Loose-patch recordings (holding voltage set to “OFF”) were performed with a new glass electrode (3–5 MΩ) filled with Ames solution. Every alpha RGC was recorded in both conditions, i.e., control and histamine. As some pDSGCs entered a depolarization block upon the addition of histamine, it was not possible to record these cells in both conditions. Therefore, some pDSGCs were recorded only under one condition, control or histamine application. For velocity tuning, cells were recorded in both conditions (before and after histamine). We did not present the faster stimuli (8, 10 or 12 Hz) for two cells that stopped responding at intermediate frequencies. For all other cells, we presented all frequencies regardless of the responses.

Intracellular current-clamp recordings from DGSCs were carried out using glass pipettes (5–9 MΩ) filled with an intracellular solution containing (in mM): 135 K Gluconate, 4 KCl, 2 NaCl, 10 HEPES, 4 EGTA, 4 ATP and 0.3 GTP (pH7.3 with KOH; Osmolarity=290 mOSm/kgH_2_O). A giga-Ohm seal was obtained before breaking in. Data were acquired at 20 kHz, and for whole-cell mode filtered at 2 kHz, with a Multiclamp 700B amplifier (Molecular Devices, CA, USA) using pCLAMP 10 recording software and a Digidata 1550 digitizer (Molecular Devices). When possible, the same cell was recorded before and after the addition of histamine.

### Visual stimuli used in patch-clamp experiments

Stimuli were generated using MATLAB and the Psychophysics Toolbox (Brainard, 1997; Pelli, 1997). A white, monochromatic organic light-emitting display (OLED-XL, 800×600 pixel resolution, 85 Hz refresh rate, eMagin, Bellevue, WA, USA) was used. The display image was projected through a 20× water-immersion objective (UMPLFLN20xW; Olympus, Tokyo, Japan), via the side port of the microscope, centered on the soma of the recorded cell, and focused on the photoreceptor layer. The diameter of the entire display on the retina was 1 mm across. The light intensity of the gray screen was 6.4x10^4^ R*rod^-1^s^-1^. For the spot stimulus, a gray background was presented for 2 s, followed by the appearance of a black (for OFF alpha RGCs) or white (for ON alpha RGCs and pDSGCs) spot on the gray background for 2 s, followed by a return to the same gray background for a further 2 s. Spots of different diameters (50-800 µm) were presented in a pseudorandom order. The total number of spikes was averaged over 5 repeats. The grating stimulus consisted of moving square-wave gratings with a spatial frequency of 400 µm. The standard temporal frequency we used was of 2 Hz (equivalent to 0.075 cycles/deg or 800 µm/s). For testing different motion velocities, we used temporal frequencies ranging from 1-12 Hz, corresponding to 400-4800 µm/s. The grating stimuli were presented in 8 different pseudorandomly chosen directions, in 45° intervals, with each presentation lasting 3 s, followed by 2.5 s of a gray screen. The stimulus was masked by a circle (diameter 400 µm) so that everything outside the circle remained gray. The total number of spikes was averaged over 3-4 repeats. For the moving-bar stimuli, a white bar (400 µm width x 900 µm length) on black background moved through the center of the screen in 8 different pseudorandomly chosen directions, in 45° intervals, at a speed of 600 µm/s. Each presentation was separated by 2 s of mean gray screen. The total number of spikes was averaged over 4 repeats.

### Data analysis of patch-clamp experiments

Electrophysiological data were analyzed offline. For loose-patch clamp recordings, spike times were extracted after filtration using a 4-pole Butterworth bandpass filter between 80 and 2000 Hz. Peri-stimulus time histograms (PSTHs) of spiking activity were calculated from 5 repeats using a bin width of 50 ms for spot stimuli. For moving grating stimuli, mean PSTHs were calculated using 25 ms bin width. For spot stimuli, the background activity was determined based on the 2 s period of initial gray screen in each trial. This provided the mean baseline activity and its SD. The bin with the highest firing rate during the spot appearance (or disappearance in the case of OFF responses in pDSGCs) was used to calculate the maximum response. Response durations were defined based on the number of all the bins during the stimulus whose value exceeded the mean baseline activity by 3 SDs (Warwick et al., 2018). Statistical tests to compare response durations were performed on log-transformed (log_10_) values of the duration, which were then distributed normally.

To analyze responses to moving gratings, we calculated the normalized vector sum as

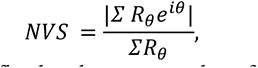

where *R_θ_* is the response in direction θ, defined as the mean number of spikes during the grating presentation (averaged over repetitions). To analyze temporal tuning, we set the NVS to 0 for cells that stopped responding (<2 spikes/s in response to gratings). We determined the PD as the direction, which most commonly had the highest response across both conditions (pre and histamine) and temporal frequencies (1-6 Hz). The normalized response in the preferred direction was calculated as the total number of spikes in the preferred direction divided by the maximum number of spikes from any temporal frequency or condition (pre and histamine). Statistical comparison was performed with a 2-way ANOVA with Bonferroni *post-hoc* correction for multiple comparisons.

In current-clamp intracellular recordings, spikes were detected as fast high-frequency events following filtration of the signal (Savitzky-Golay filter, 1^st^ order). To analyze the effects of histamine on the spontaneous activity of pDSGCs, we calculated the firing rate after histamine reached the bath and compared it to the mean firing rate of the baseline (calculated from a 50-second-long time window after adding histamine or in Ames medium). To analyze the spike shape, spikes were aligned to their peak, averaged for each recorded cell and plotted in a time window that started 100 ms prior to the peak and ended 40 ms after the peak. The spike half-width was quantified as the duration where the normalized voltage was more than half the spike’s amplitude. The baseline voltage was calculated as the average voltage in the first 20 ms in the aligned spikes for each cell.

### Two-photon Ca^2+^ imaging

Two-photon Ca^2+^ imaging (Bruker microscope equipped with a Spectra-Physics Mai-Tai laser) from the ganglion cell layer of the isolated retina of mice expressing GCaMP6f in RGCs (Thy1-GCaMP6f) was carried out on an area of 140x140 µm^2^ at 6 Hz. For the UV stimuli, a modified projector (M109s DELL, Austin, TX, USA) containing a UV LED (NC4U134A, peak wavelength 385 nm; Nichia, Anan, Japan) was used (Borghuis et al., 2013). The image was projected onto the retina via the microscope’s condenser and created on the photoreceptors layer using two converging lenses (LA4372, LA4052; Thorlabs). The field of view was positioned in the center of the visual stimulus. In the histamine studies, control experiments, in which histamine was not added, were performed using the same time course as the histamine experiments. Histamine was added 10 min prior to imaging under histamine conditions. Histamine was washed out with Ames’ solution for 45 mins prior to imaging.

### Visual stimuli used in two-photon Ca^2+^ imaging experiments

For UV spot stimuli, a spot (300 µm in diameter) of increased luminance (2.8x10^4^ R*rod^-1^s^-1^) centered on the field of view (140 x 140 µm) appeared for 2 s. The ΔF/F was averaged over 3 trials. The moving grating and bar stimuli had the same properties as those used for the patch-clamp experiments, except that they were of UV light and 3 repeats were carried out.

### Data analysis of two-photon Ca^2+^ imaging experiments

ROIs were manually selected using an average projection of the responses to the moving bar stimulus (all directions and repeats) with ImageJ software (Schneider et al., 2012). Each field of view contained between 9-38 RGCs. To determine whether an RGC responded to the histamine application, the mean baseline and SD were calculated from the 30 s immediately prior to histamine arriving in the bath. RGCs that exceeded 6 SDs over the mean baseline in the 40 s period after histamine’s arrival were counted as responsive to histamine. Prior to any visual stimulus, the RGC layer was imaged for 30 s. The latter 15 s of this pre-stimulus were taken as the baseline and used to calculate the ΔF/F. To determine whether an RGC was responsive to a spot stimulus, a threshold of 3 SDs above the mean baseline was set during the ON period (appearance of white spot) and OFF period (2 s after the spot disappeared). An additional threshold was set 3 SDs below the mean baseline during the ON period. Any RGY whose ΔF/F trace crossed any of these thresholds was counted as light responsive. To determine whether an RGC response polarity was ON or OFF, we used an ON-OFF index (OOI): 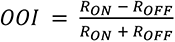, where R is the maximum amplitude (ΔF/F). RGCs with a negative OOI were deemed OFF RGCs, whereas those with a positive OOI were deemed ON RGCs. As the responses of some RGCs changed with time, only RGCs that had the same response polarity (ON, OFF or non-responsive) under the pre and histamine-washout conditions were included (54% (276/512) for histamine data set; 70% (98/140) for control data set). In the statistical analysis using Fisher’s exact test, all changing RGCs were grouped together and all non-changing RGCs were grouped. RGCs were classified as changing if they lost, gained or changed response polarity. Response amplitudes before and after histamine were calculated in units of SDs (based on the 15 s baseline recording) and were plotted against each other, and the absolute distance from the unity line was calculated and their distribution between the control and histamine datasets were compared using a Kolmogorov-Smirnov test. In Figures S4A-C, RGCs were further divided into transient and sustained groups. For OFF RGCs, this was done by calculating a transient-sustained index where the mean of the response trace (during the OFF period) was divided by the maximum amplitude (during the OFF period). Those RGCs with a transient-sustained index > 0.4 were deemed sustained, whereas those < 0.4 were considered transient. For ON RGCs, the transient-sustained index was calculated in the same way, except for using the ON period and a further step where, if the maximum amplitude occurred during the first half of the ON period, the transient-sustained index was further divided by 2. Those RGCs with a transient-sustained index > 0.35 were deemed sustained, whereas those < 0.35 were considered transient. To determine whether an RGC was direction-selective in response to moving gratings, we computed the normalized vector sum: 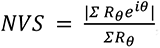, where *R_θ_* is the area under the curve in direction *θ*. RGCs with an NVS>0.25 were counted as DS. Of these RGCs, those that reduced their NVS by more than 0.12 upon the histamine application were deemed to have reduced direction-selectivity.

### Multi-electrode array recordings

Multi-electrode array (MEA) recordings were performed on isolated retina using multi-electrode arrays of 252 electrodes (MultiChannel Systems, 252 electrodes, 30 µm diameter, 100 µm minimal electrode distance). The retina was mounted on the MEA with the ganglion cell layer facing down. The MEA was placed in the head stage with constant perfusion of oxygenated bicarbonate-buffered Ames medium at a flow rate of 3.5 ml/min; a heating pad placed below the array maintained the temperature at 33.2°C. Data acquisition started one hour after the retina was placed in the chamber, to let the retina adapt. Extracellular voltage signals were amplified and digitized at 20 kHz and saved for offline analysis.

Visual stimuli were created in MATLAB (version R2018a), using Psychophysics Toolbox (Brainard, 1997; Pelli, 1997) and a custom GUI, and were projected via a monochromatic white OLED display (eMagin, EMA-100309-01 SVGA+, 600x800 pixels, 60Hz refresh rate) through a telecentric lens (Edmund Optics, 2.0X, #58-431) onto the photoreceptors. The pixel size on the retina was 7.5 µm. At maximum brightness, the irradiance used in the experiments was 2.6 µW/cm^2^, resulting in 2.43x10^4^ mouse rod isomerizations (R*rod^−1^s^−1^), whereas the minimum brightness was 7.04x10^1^ R*rod^−1^s^−1^.

### Visual stimuli for MEA recordings

We used a battery of visual stimuli in the MEA recordings. All stimuli were presented in full-field, covering the entire electrode array (electrode area: 1500x1500 µm, stimulus size was always at least 2250 µm in diameter). We recorded 30 s of spontaneous baseline activity before presenting each stimulus. Stimuli were repeated 5 times unless otherwise specified. The full-field stimulus sequence was 3 s black, 2 s white, 3 s black, and illuminated the full screen in uniform intensity. Moving square-wave gratings of 100% contrast with a spatial frequency of 397.5 µm (0.075 cycles/deg) moved in 8 directions, in 45° intervals, in a randomized order with a speed of 795 µm/sec (2 Hz). Each moving grating was presented for 4 s, preceded and followed by 2 s of mean gray background intensity. To investigate temporal tuning we presented the same stimulus at temporal frequencies of 1, 2, 4, 6, 8, 10 and 12 Hz in a random order (3 s grating, 4 repeats).

### Pharmacology in MEA experiments

We carried out MEA recordings to determine the effective histamine concentration (see *Histamine concentration calibration*) and to establish the specific effects of histamine application and chemogenetic activation of retinopetal axons by the synthetic ligand clozapine-N-oxide (CNO) on RGCs’ output. We recorded three runs of the same set of visual stimuli in the pre (no drug added), with histamine/CNO and after-histamine-washout conditions. In experiments where histamine/CNO were added to the bath, we recorded the spontaneous activity of RGCs in darkness (OLED switched off) for 10 min before switching to histamine (5 or 10 µM)/CNO (15 µM). We recorded a TTL pulse when the switch to Ames+drug occurred. Histamine/CNO was washed in for at least 10 min before we displayed the visual stimuli again. The washout was performed for 1 h. For the histamine studies, control experiments were performed with the same time course as the histamine experiments, but without washing in histamine (the time points are indicated as run 1, run 2 and run 3 in the Figures and captions). For the chemogenetic experiments, CNO was added to both control (non-injected HDC-Cre mice) and hM3Dq-AAV-injected mice retinas. The histamine solution was prepared as described in *Histamine application and pharmacology*, while CNO (Clozapine N-oxide dihydrochloride, Tocris, Cat. No. 6329) was dissolved in water to make a stock solution of 20 mM (aliquots stored at -20°C) and then further diluted in Ames solution to reach a final concentration of 15 µM.

### Histamine concentration calibration

In order to obtain a dose-response curve for different concentrations of histamine, we performed MEA experiments in which we successively washed in 1, 2, 5, 10 and 20 µM of histamine (prepared as previously described) to the bath solution while recording the baseline activity of RGCs in darkness (OLED switched off). Each concentration was washed in for approximately 2 min before switching to the next concentration. We recorded a TTL pulse whenever we switched to the next concentration. Control experiments were performed in the same way without washing in any histamine. The time points t_1_-t_5_ in the control experiments, shown in Figure S2, correspond to histamine concentrations 1, 2, 5, 10 and 20 µM, respectively.

### Data analysis of MEA experiments

Spike sorting was performed using Kilosort2.0 (Pachitariu et al., 2016, 2018), with subsequent manual curation in Phy (Rossant and Harris, 2013; Rossant et al., 2016). We only included well-separated units in our analysis. Data were analyzed using custom-written scripts in MATLAB (version R2018b and R2019b).

### Histamine concentration calibration analysis

To analyze the concentration-dependent effect of histamine, spike times were binned using time bins of 1 s. We calculated the mean firing rate over a window of 30 s just before the switch to the next concentration occurred (gray shaded bars in Figure S2A). We then calculated the difference between each mean firing rate to the baseline (0 µM histamine or t_0_ in control experiments, Figures S2C and S2D). The percentage of responsive RGCs for each concentration (Figure S2E) was determined as described later (*Histamine wash-in analysis)*. In total, 154 RGCs from 3 retinas were used in the histamine calibration experiments, and 361 RGCs from 4 retinas in the control experiments.

### Histamine wash-in analysis

To analyze the effects of histamine on the spontaneous activity of RGCs recorded using the MEA, we calculated the firing rate over the duration of the wash-in (15-20 min) using a bin width of 1 s. We focused on the time window of 60-300 s (4 min duration) after histamine reached the bath and compared the mean firing rate of RGCs to their baseline firing rate (calculated from a 60 s time window in Ames medium just before histamine was added). We only included RGCs with a minimum firing rate of 1 Hz across the duration of the wash-in (n=640/763 from 7 retinas for control, and 712/765 RGCs from 6 experiments for histamine). The magnitude of change in spontaneous activity was calculated as the difference in firing rate from that of the baseline. We defined cells as responsive to histamine if they crossed an upper or lower threshold, determined by the upper and lower 2.5% quantile of the control distribution (3.5 and -4.6 spikes/s, respectively, Figure 2B, red vertical lines). We used the same thresholds to determine the percentage of responsive RGCs in the concentration calibration experiments (Figure S2E). The percentage of RGCs with increased firing rates upon histamine application was compared to the control data set using a two-sample t-test for unequal variances.

### Analysis of MEA light responses

To classify RGCs as ON, OFF or ON-OFF, we defined an ON-OFF index (OOI), calculated from the response to the full-field stimulus:

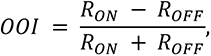

where R_ON_ and R_OFF_ are the spike counts during 2 s of light ON or OFF, respectively. This results in values in the range of [-1, 1], where ON RGCs will have a positive OOI, OFF RGCs a negative OOI, and ON-OFF RGCs will have OOIs in between, depending on whether their ON or OFF response is more prominent. The PSTH in response to the spot stimulus (Figure 2C) was calculated as the mean PSTH over 5 repetitions using a bin width of 50 ms.

To identify direction-selective ganglion cells (DSGCs), we analyzed the response to moving gratings. Prior to analysis, motion directions were aligned to retinal coordinates. We then calculated the normalized vector sum as above (*Data analysis of Patch clamp recordings)*.

The preferred direction (PD) was defined as the angle of the vector sum. The direction-selectivity index (DSI) was calculated as

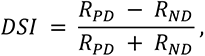

where *R*_PD_ and *R*_ND_ are the responses in the direction closest to the PD and the one opposite to it, respectively. Similarly, we calculated an orientation-selectivity index as

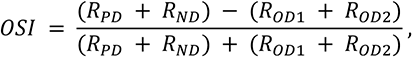

where *R*_PD_ and *R*_ND_ make up the response in the preferred axis, and R_OD1_ and R_OD2_ are the responses in both orthogonal directions. We only considered RGCs that had a mean firing rate > 1 Hz during the grating stimulus. Cells with a NVS > 0.15, a DSI > 0.3 and OSI < 0.3 were considered as DSGCs. We obtained 39/591 (5 retinas) and 43/485 DSGCs (4 retinas) for the control and histamine experiments, respectively.

To study the effects of histamine on DSGCs, we only included DSGCs that displayed reversible changes in their evoked responses to moving gratings, based on the assumption that gradual changes over time cannot necessarily be attributed to the effect of histamine (although this strict criterion might have skipped long-lasting effects mediated by the activation of histamine G-protein coupled receptors). Indeed, we found that also in control experiments, some DSGCs displayed gradual changes over time (>50% of cells gradually increased their evoked responses over time; data not shown). Therefore, we only considered DSGCs that reversibly changed in the DSGC analysis. DSGCs were considered to reversibly change if their evoked response to moving gratings either increased or decreased in at least 4 directions by at least 1 SD (calculated over 5 repetitions in the pre condition), and showed a washout effect of at least 30% of the initial change. We then determined the proportion of reversibly changing DSGCs that decreased or increased their directional tuning upon histamine application based on the NVS. To assess the effects of histamine application on directional tuning, we performed a Friedman test on the NVS of all the DSGCs with reversible changes in their light responses (n=18) and then performed subsequent pairwise comparisons using a Wilcoxon signed-rank test.

To analyze responses of DSGCs to moving gratings of different temporal frequencies, we classified RGCs as DSGCs based on their response to 2 Hz moving gratings, using the same criteria as before. We obtained 27/392 DSGCs (3 retinas). We defined the PD as the direction closest to the angle of the NVS for gratings moving at 2 Hz. We then performed a fast Fourier transform (FFT) on the PSTHs of each DSGC’s PD (mean PSTH over 4 repetitions, using 25 ms bin width, with mean subtracted) for all temporal frequencies presented and calculated the amplitude of the first harmonic at each stimulus frequency. Statistical comparison was performed with a 2-way ANOVA with Bonferroni *post-hoc* correction for multiple comparisons.

### Analysis of DREADD experiment

To determine the effects of chemogenetic activation of retinopetal axons on the spontaneous activity of retinal RGCs recorded on the MEA, we calculated the firing rate over the duration of the wash-in (10 min) using a bin width of 50 ms. In particular, we focused on a 5-min-long time window after CNO reached the bath (starting 60 s after the drug-TTL pulse was sent) and compared the mean firing rate of RGCs to their baseline firing rate (pre, mean over 60 s before the drug-TTL pulse was sent). We only included RGCs with a minimum firing rate of 1 Hz (660/717 RGCs from 5 retinas (5 mice) in control and 1172/1290 RGCs from 13 retinas (9 mice) in hM3Dq experiments, respectively). Right and left retinas from the same mouse were recorded only when mice were injected bilaterally in the TMN; otherwise, the contralateral dorsal retina was recorded. The magnitude of change in spontaneous activity, recorded in darkness, was calculated as a change in the firing rate from the pre to the CNO condition, and the distributions in the control and injected retinas were compared using a Kolmogorov-Smirnov test. We defined RGCs as responsive to CNO application in hM3Dq retinas if they crossed an upper or lower threshold, determined by the upper and lower 2.5% quantile of the distribution obtained in the control experiments (7.11 and -1.87 spikes/s, respectively). The percentage of RGCs that changed their activity upon CNO application was compared to the control data set using a binomial test. To check whether CNO’s effect was time-dependent, we calculated the mean firing rate in three 60-sec-long windows after 1.5, 4.5 and 6.5 min (t_1_, t_2_ and t_3_) from the drug wash-in and compared it to the baseline (Figure S8C). To study the effects of activation of histaminergic retinopetal axons on DSGCs, we only included RGCs that displayed reversible changes in their evoked responses to moving gratings (as in *Analysis of MEA light responses).* The gratings stimulus was presented to 5 and 11 retinas in control and hM3Dq experiments, obtaining in total 34/698 and 30/1043 DSGCs, respectively.

### Human experiments

The study protocol was reviewed and approved by the Institutional Helsinki Committee of Kaplan Medical Center, Rehovot, Israel. Nine healthy men and women between the ages of 18-50 were recruited. All recruited subjects provided signed informed consent. Exclusion criteria included chronic disease, taking regular medications, taking medication in the two weeks prior to the study visit, eye diseases that affect the functions of the optic nerve or the retina, farsightedness or family history of narrow-angle glaucoma, difficulty urinating or known enlargement of the prostate, pregnancy and breastfeeding. One patient was excluded from the study due to an inability to complete the study tests in light of known ADHD.

In this clinical, crossover, single-blind trial, each volunteer participated in two visits, at least two weeks apart. During one visit, 2 mg of dimetindene maleate (Fenistil, Glaxosmithkline) oral drops (1 mg/ml) diluted in 200 ml of sweetened water was given, and during the other visit a placebo was given. Dimetindene maleate was chosen based on safety (commonly used as an anti-allergy medication) and because it has been reported to cross the blood brain barrier and hence would likely reach the retina (Mahdy and Webster, 2017; Noguchi et al., 1992). The order of the visits was random. The placebo or drug were taken between 8:00 AM and 12:30 PM; tests to assess visual function were performed two hours after dimetindene maleate/placebo consumption. Refraction and Best Corrected Visual Acuity were measured in both eyes separately at the beginning of each visit, using a Snellen chart, based on autorefractometer results and an optometrist exam.

During each visit, we used a Heidelberg Spectralis device to perform macular Optical Coherence Tomography (OCT), OCT retinal nerve fiber layer (RNFL) and OCT angiography (OCTA) to ensure that Fenistil does not cause any morphological changes in the retina or the blood vessel density in it. We used the macular OCT and OCT RNFL images to compare the Central Macular Thickness (CMT) and mean RNFL thickness, respectively. To quantify retinal vascular density, the en-face images of different vascular retinal layers obtained with OCTA were processed by ImageJ 1.52v software. Images were binarized according to Niblack’s method and a grayscale mean was calculated and transformed to coverage in percent, after subtracting the Foveal Avascular Zone (Fujiwara et al., 2017). No differences were observed between the placebo and H_1_R antagonist (Table S2).

Visual field tests were carried out using the Humphrey visual field analyzer (HFA). The central visual field was evaluated using the HFA 10-2 program of automated perimetry with the Swedish Interactive Threshold Algorithm (SITA) standard strategy, and the peripheral visual field was assessed using the HFA 60-4 program of automated perimetry with the SITA standard strategy. Perimetry, which refers to the systematic measurement of the visual field, measures sensitivity to stimuli at multiple locations in the visual field while monitoring fixation. Each eye is tested separately. We used a white-on-white size III (0.4 mm) target with a background luminance of 31.5 apostilbs (asb) in all the tests. The Humphrey perimeter tests light intensities over 5 orders of magnitudes, from 10,000 asb to 0.1 asb. Every log order change in light intensity corresponds to 10 dB, such that the machine can measure sensitivities over a 50 dB range. The SITA developed for the Humphrey perimeter estimates threshold values for each point of the visual field based on responses to stimuli presented at that location, as well as information gathered from nearby locations. In the 10-2 perimetry, 68 points in the central 10 degrees of the visual field were measured, including the foveal sensitivity. In the 60-4 perimetry, 60 points were measured between 30 to 60 degrees of the visual field. Fixation is assured by mapping the blind spot, and then retesting the blind spot throughout the visual field test. Positive responses during retesting of the blind spot are assumed to reflect loss of fixation; the ratio between fixation losses and the number of times the blind spot was tested was less than 20% in all tests.

### Statistical analysis

Statistical parameters, including the exact value of n, mean ± standard deviation (SD) and statistical significance, are reported in the text and figure legends. The cutoff for significance was *p*<0.05, and the significance level is marked by \**p*<0.05, \*\**p*<0.01, \*\*\**p*<0.001, and \*\*\*\**p*<0.0001. Statistical analysis was performed with MATLAB (Natick, Massachusetts: The MathWorks Inc) or Prism (Graphpad, San Diego, CA, USA). Welch’s one-way ANOVA (MATLAB Central File Exchange., 2021) was performed with MATLAB.

### Data availability

The data that support the findings of this study are available upon request.

