## Supplemental Information for "Top-down modulation of the retinal code via histaminergic neurons of the hypothalamus"

### Supplementary Figure 1

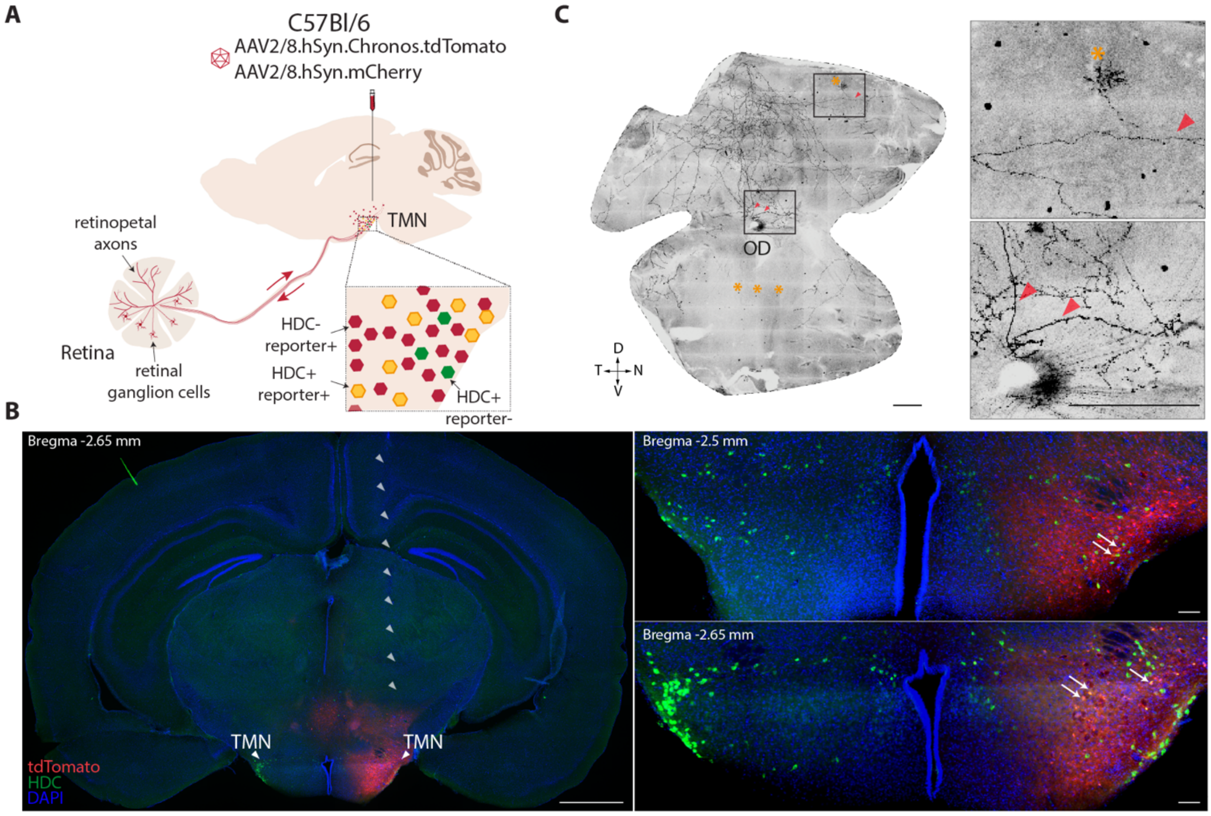


**Figure S1. Retinopetal projections in WT mice.** (**A**) Schematic of the microinjection of viral tracers (AAV2/8.hSyn.Chronos.tdTomato or AAV2/8.hSyn.mCherry) into the TMN region of C57Bl/6 mice. Brains and retinas were analyzed after 4-6 weeks. The virus infected both $HDC^{+}$ and $HDC^{-}$ cells, and the presence of retinopetal axons was often accompanied by labeled cell bodies in the retina. (**B**) Left: Whole-section fluorescent photomicrograph of a C57BL/6J mouse brain slice following a unilateral injection of AAV2/8.hSyn.Chronos.tdTomato into the TMN. Red, tdTomato expression; green, immunostaining for anti-HDC; blue, DAPI. Right: Two serial coronal sections of the TMN displaying expression of double-labeled cells (yellow), indicated by white arrows. The distance from Bregma is noted on the left top corner, in mm. (**C**) Left: Tiled fluorescent images showing retinopetal axons in an example retina 4 weeks after injecting AAV8.hSyn.mCherry into the TMN. Right: Higher magnification of the regions indicated by the black boxes. Red arrowheads indicate axonal branches running through the dorsal retina. Orange asterisks (*) mark positive cell bodies. Scale bars, 1000 µm for (b, left), 100 µm for (b, right) and 500 µm for (c). Abbreviations: TMN – tuberomammillary nucleus, OD – optic disc, D, N, T, V – dorsal, nasal, temporal, ventral.

### Supplementary Figure 2

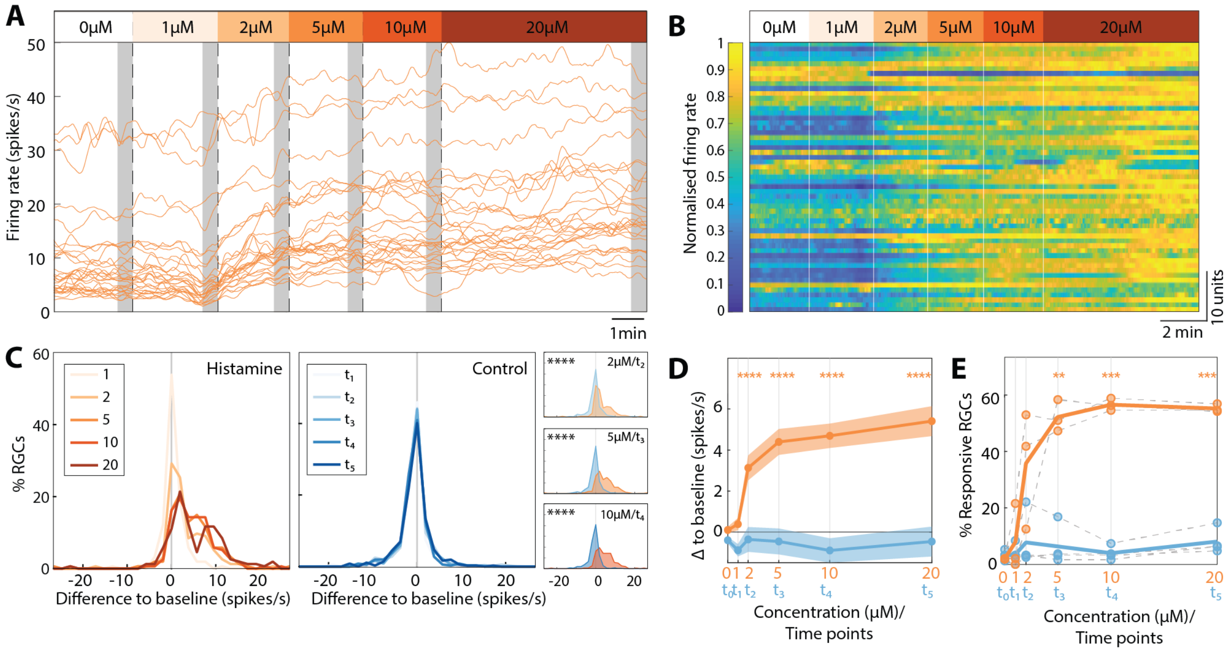


**Figure S2. Concentration-dependent effects of histamine.** (**A**) Baseline firing rate of 30 example RGCs recorded from a dark-adapted retina on the MEA to the wash-in of different concentrations of histamine (1, 2, 5, 10, 20 μM; shown above). Gray shaded areas mark the time when the final firing rates for each concentration were determined (30 s duration before washing in the next concentration). (**B**) Normalized firing rates of all the RGCs from the same experiment shown in (A) (n=55 cells). (**C**) The difference in the firing rate distribution between each concentration (from gray area in (A)) and the baseline in the histamine (orange, left) and control (no histamine was added; blue, middle) experiments. The histamine wash-in increased the firing rates for concentrations above 2 μM. Right/inset: Overlaid distributions for control and histamine for concentrations/time points of 2 μM/$t_2$, 5 μM /$t_3$ and 10 μM /$t_4$ (Kolmogorov-Smirnov test). (**D**) Mean±SEM dose-response curves of the change in the firing rate for histamine (orange, n=154 units) and control (blue, n=361 units). The statistical comparison was to the baseline concentration (0 μM/$t_0$). Significances are displayed above only for significant increases (one-tailed paired t-test with Bonferroni-correction). (**E**) The percentage of responsive units to each concentration/time point of the histamine/control (determined as in Figure 2) single experiments (gray dashed lines) and mean across experiments for histamine (orange bold line, n=3 retinas) and control (blue bold line, n=4 retinas). Statistical notation as in (D) (one-tailed paired t-test). Note that although we did not add any histamine in the control experiments, we followed the same time course as in the histamine experiments. Timepoints $t_1$-$t_5$ in (C) correspond to histamine concentrations 1, 2, 5, 10, 20 μM. *: $p$<0.05, **: $p$<0.01, ***: $p$<0.001. ****: $p$<0.0001.

### Supplementary Figure 3

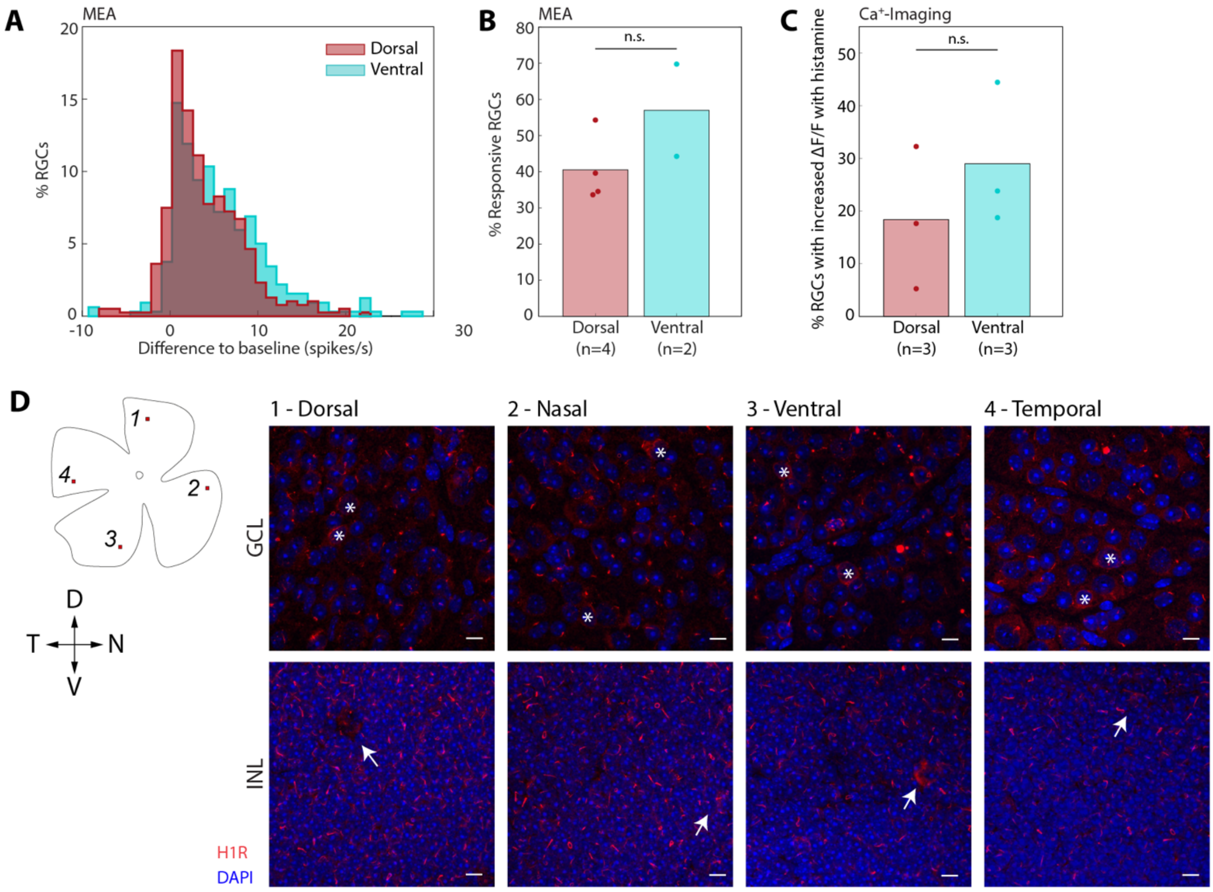


**Figure S3. Histamine affects RGCs in both dorsal and ventral parts of the retina.** (**A**) Difference in RGC firing rates compared to those of the baseline upon histamine wash-in (recorded on the MEA), separated into dorsal (n=387 RGCs from 4 retinas) and ventral (n=319 RGCs from 2 retinas) retina regions ($p$=0.0003, Kolmogorov-Smirnov test). (**B**) Mean percentages of RGCs in which histamine increased the baseline firing rate, separated into dorsal and ventral retina regions, with data points of single experiments overlaid (n=4 and 2 retinas for dorsal and ventral, respectively; *n.s.*, independent-sample t-test). (**C**) Like (b), but for $Ca^{+}$-imaging data (n=3 and 3 retinas for dorsal and ventral, respectively; *n.s.*, independent-sample t-test). (**D**) Retinal wholemount immunostained for histamine 1 receptor ($H_1R$; red). Images, shown as maximum intensity projections, of four different areas (1-4, inset on the left) in the retina at the level of GCL (top) and INL (bottom) focal planes, respectively. Positive puncta were found in all locations in the retina. A subpopulation of $H_1R$-positive cell bodies was found in the INL (white arrows) and GCL (white asterisks). Scale bar, 10 µm.

### Supplementary Figure 4

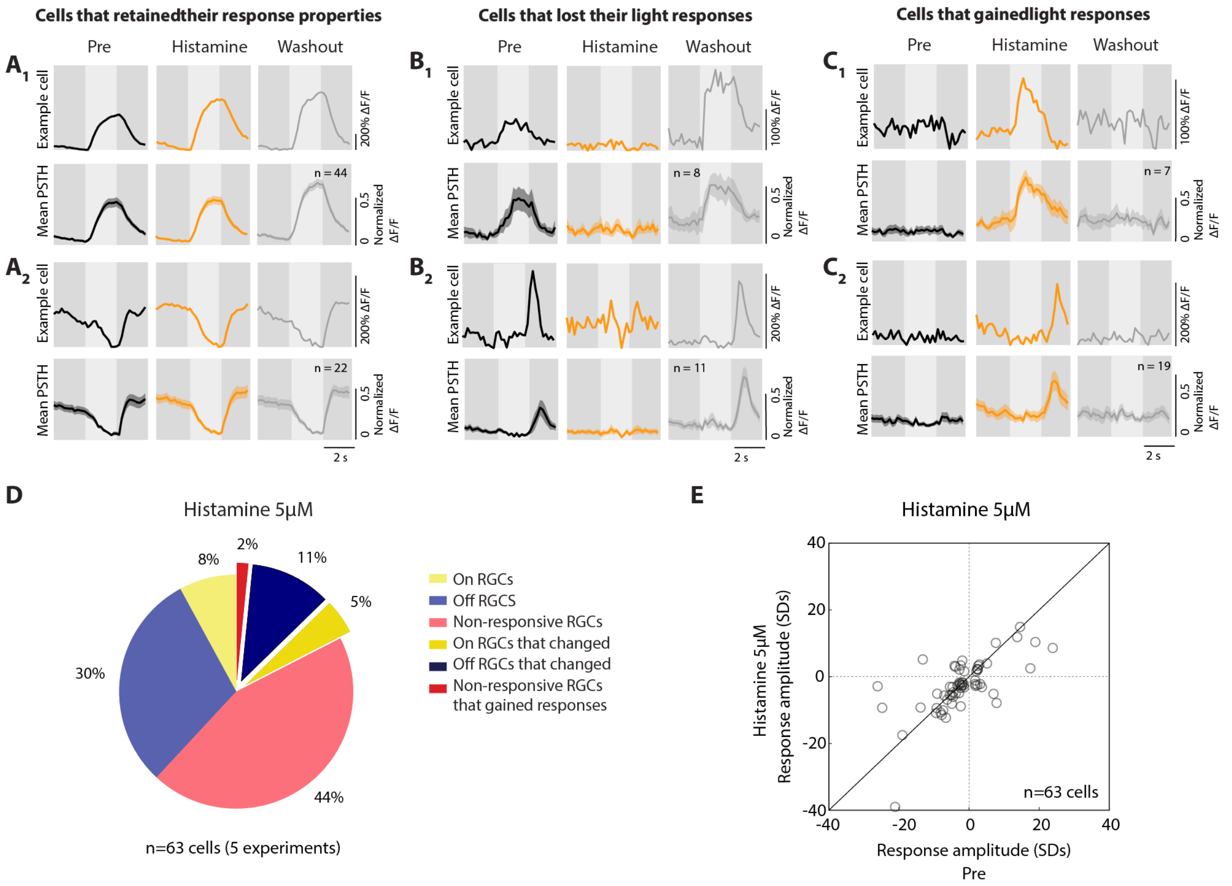


**Figure S4. Two-photon $Ca^{2+}$ imaging reveals that histamine affects RGC light responses to a spot stimulus.** (**A**) Example traces (above) and mean PSTHs (below) of sustained-ON RGCs ($A_1$) and sustained-OFF RGCs ($A_2$) that retained their light responses with histamine. (**B**) Example traces (above) and mean PSTHs (below) of sustained-ON RGCs ($B_1$) and transient-OFF RGCs ($B_2$) that reversibly lost their light responses with histamine. (**C**) Example traces (above) and mean PSTHs (below) of non-responsive RGCs that either gained a sustained-ON (i) or transient-OFF response (ii). (**D**) Pie chart showing the percentages of RGCs that changed their polarity with 5 µM histamine (17.5%, 11/63 RGCs; $p$=0.0028 compared with control data set (Figure 3B), Fisher's exact test). (**E**) Population data showing how response amplitudes to the spot stimulus change with 5 µM histamine. ON RGCs (ON-OFF index > 0) are plotted as having a positive response amplitude. OFF RGCs (ON-OFF index < 0) are plotted as having a negative response amplitude. (**A-C**) RGCs responses were categorized as ON or OFF based on an ON-OFF index (see *Methods*), and as either transient or sustained based on a transient-sustained index specific to the RGCs polarity (see *Methods*).

### Supplementary Figure 5

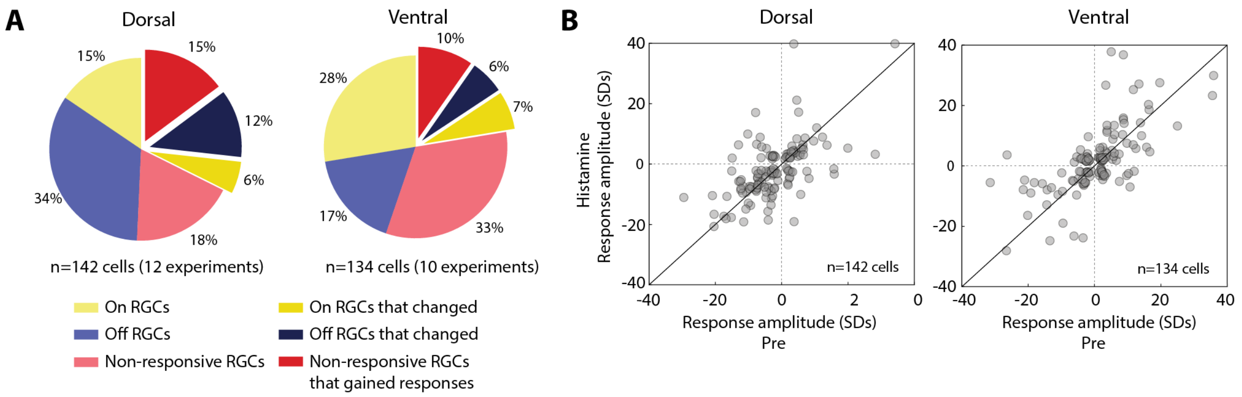


**Figure S5. Histamine affects RGC light responses in both dorsal and ventral parts of the retina.** (**A**) Pie charts showing the percentages of RGCs that retained and changed their light responses to spot stimulus with histamine (20 µM) in the dorsal and ventral retina in the two-photon $Ca^{2+}$ imaging experiments. In the dorsal retina, 46/142 (32.4%) RGCs changed their polarity preference compared to 30/134 (22.4%) RGCs in the ventral retina ($p$=0.079, *n.s.,* Fisher's exact test). (**B**) Change in the response amplitude with histamine in dorsal and ventral retinas. Responses of ON RGCs (ON-OFF index > 0) are plotted as positive amplitudes, and responses of OFF RGCs (ON-OFF index < 0) as negative response amplitudes. The absolute distance from the unity line did not differ between the dorsal and ventral populations ($p$=0.1323, *n.s.*, Kolmogorov-Smirnov test).

### Supplementary Figure 6

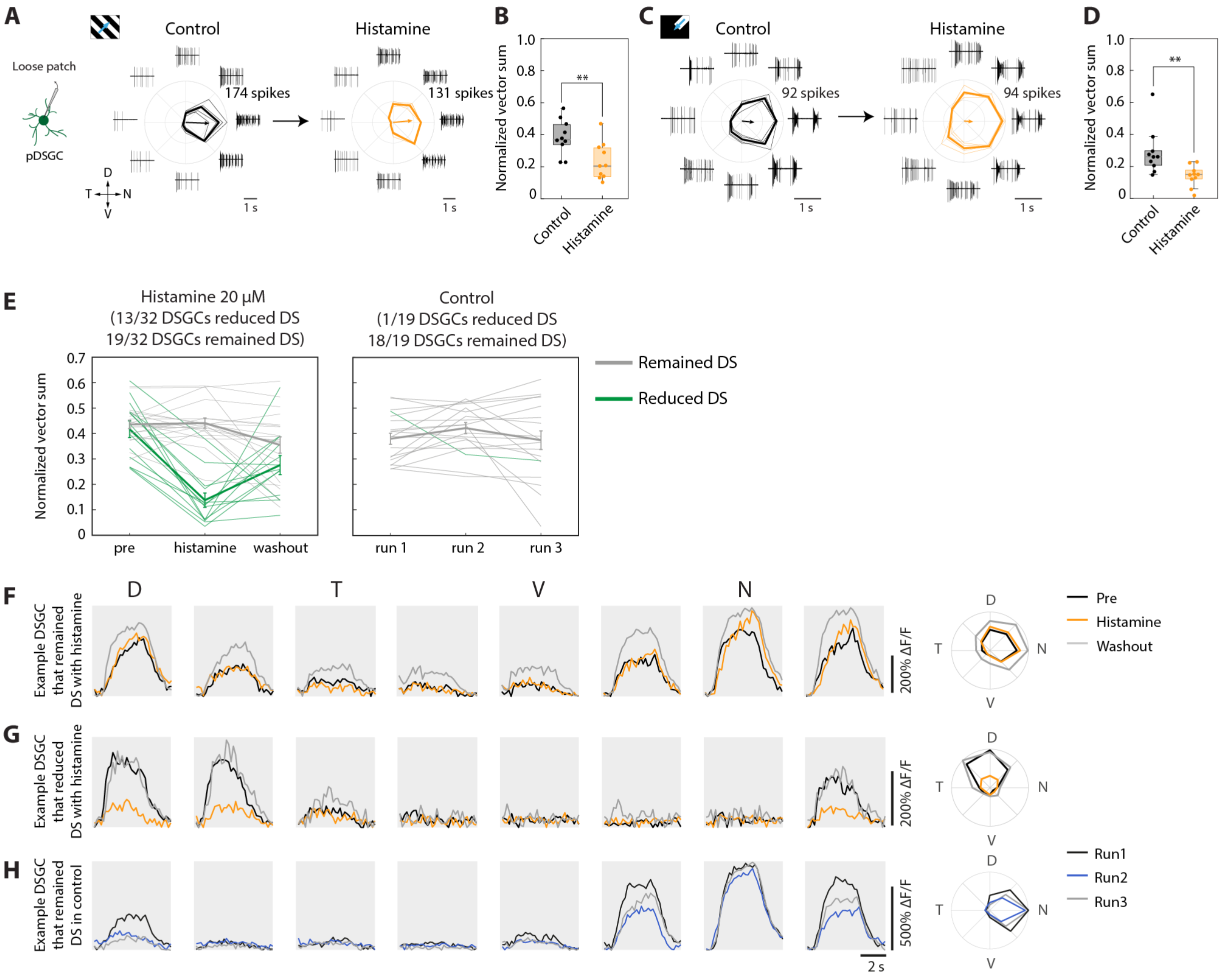


**Figure S6. Two-photon Ca$^{+2}$ imaging reveals that histamine reduces direction selectivity.** (**A**) Responses of an example pDSGC to moving gratings (2 Hz, 800 μm/s) in control and after washing in histamine (20 μM). Polar plots represent the mean response (bold line); thin lines show single repetitions. Arrows represent the normalized vector sum (NVS). Example traces illustrate the response data to each of the 8 directions. (**B**) Population data of the NVS to moving gratings ($p$=0.0086, independent-sample t-test; n=10 pDSGCs in each group). (**C, D**) Like (A, B), but for moving bars (600 μm/s; $p$=0.0083, independent-sample t-test). (**E**) Normalized vector sum (NVS) of DSGCs (DS; NVS>0.25) when presented with moving gratings. DSGCs marked in green reduced their NVS by more than 0.12. In the histamine data set (left), 13/32 DSGCs had reduced NVS compared with 1/19 DSGCs in the control data set (right) ($p$=0.0084, Fisher's exact test). Bold lines indicate the mean±SEM. (**F**) Mean trace of an example DSGC that remained DS with histamine when presented with gratings moving in 8 different directions. Far right: Polar plot showing the DSGC's responses to the 8 directions in the pre (black), histamine (orange) and washout (grey) conditions. (**G**) Same as in (F) for a DSGC that reduced its NVS with histamine. (**H**) Same as in (F) for a DSGC in the control data set.

### Supplementary Figure 7

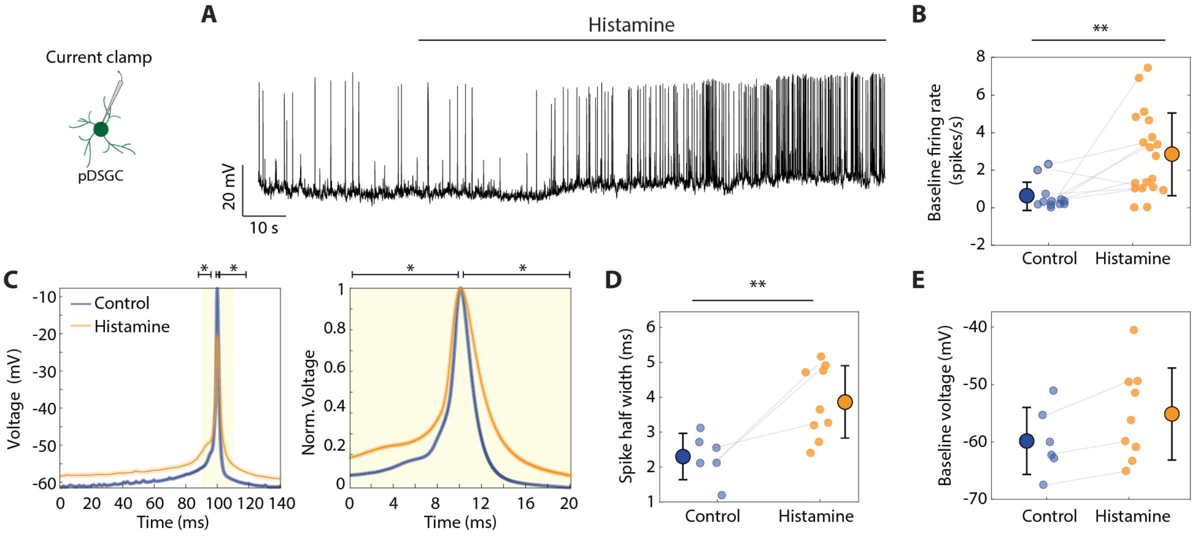


**Figure S7. Intracellular recordings to characterize the effect of histamine on pDSGCs.** (**A**) Current clamp recording from a pDSGC during histamine perfusion (5 µM, horizontal black line above). (**B**) Comparison of the mean firing rate (averaged over 50 seconds) before and after the histamine application for pDSGCs ($p$=0.0018, control: 0.61±0.75 spikes/s, n=12; histamine: 2.84±2.20 spikes/s, n=19). Gray lines indicate paired cells. (**C**) Spike shape (left) and normalized spike shape (right, close-up of the yellow shaded region in the left graph) of pDSGCs (mean±SEM) for control (n=6) and histamine (n=9). (**D, E**) Mean spike half-widths calculated from the normalized spike shape (D; $p$=0.0048, control: 2.31±0.66 ms; histamine: 3.87±1.04 ms) and baseline membrane potential (E; *n.s.*, control: -59.81±5.83 mV; histamine: -55.12±8.00 mV) in control and in the presence of histamine (control n=6; histamine n=9). Gray lines indicate paired cells. Values represent mean±SD (indicated on the side) of recorded cells unless specified otherwise. Comparisons were performed with independent-sample t-test. *: $p$<0.05, **: $p$<0.01.

### Supplementary Figure 8

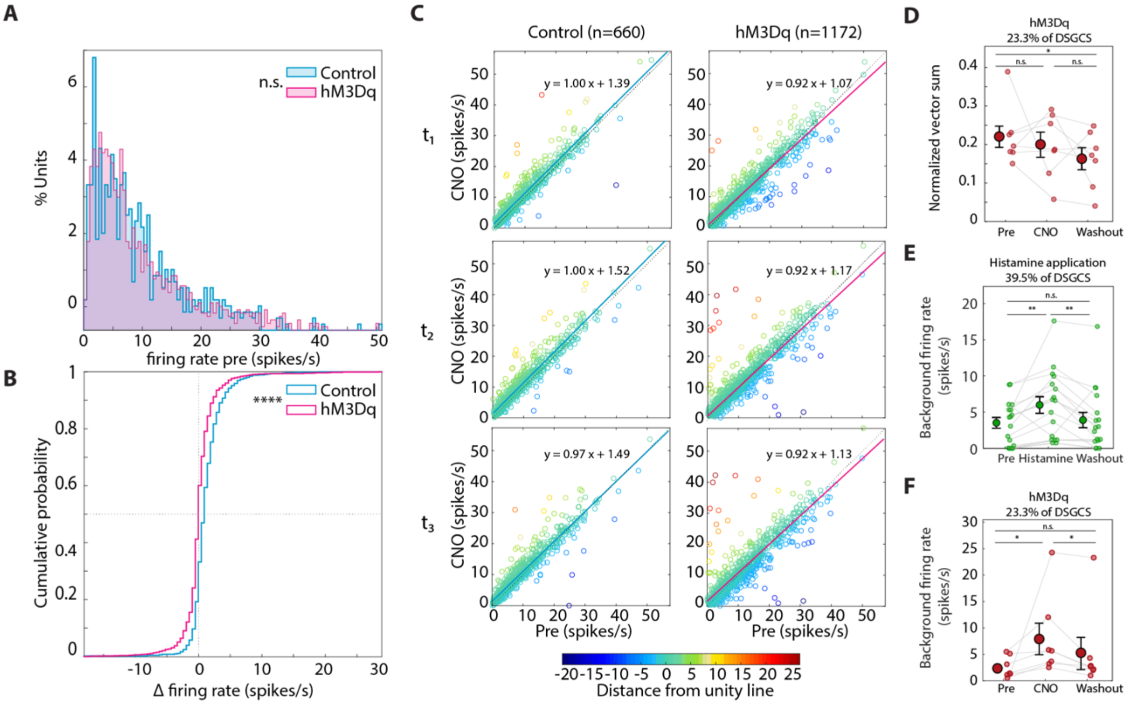


**Figure S8. Activating histaminergic retinopetal axons modulates RGC activity.** (**A**) Distribution of the firing rate before CNO administration for the control (non-injected mice, cyan) and hM3Dq (pink) data sets. Distributions are not significantly different (*n.s.*, Wilcoxon signed-rank test). (**B**) The cumulative density function of the change in firing rate (CNO-pre) for the control (cyan) and hM3Dq (pink) data sets. The hM3Dq RGC population is significantly skewed toward the left ($p \ll 0.0001$, Kolmogorov-Smirnov test). (**C**) Comparison of the firing rates before (x-axis) and after (y-axis) CNO application in the control (top) and hM3Dq (bottom) data sets. Timepoints $t_1$-$t_3$ correspond to 60-sec-long time windows 1.5, 4.5 and 6.5 min after the drug wash-in, respectively. Individual cells are color-coded according to their distance from the unity line. (**D**) The NVS of all the DSGCs with reversibly changing light responses to gratings (from Figure 5I (red), n=7 cells) with mean±SEM overlaid (black) (*n.s.* (pre - CNO), *n.s.* (CNO - washout), $p$=0.046 (pre - washout), Wilcoxon signed-rank test). Mean±SEM: 0.22±0.03, 0.19±0.03, 0.16±0.03 in pre, CNO and washout, respectively. (**E**) Background firing rate at mean gray intensity for all DSGCs that decreased their tuning with histamine (n=17 cells, green cells from (Figure 5E)) calculated for all conditions before the grating stimulus was presented ($p$=0.0036 (pre-histamine), $p$=0.0065 (histamine-washout), *n.s.* (pre-washout), Wilcoxon signed-rank test). (**F**) The baseline firing rate during the pre-stimulus (mean gray) of all the DSGCs (from Figure 5I (red), n=7 cells, $p$=0.016 (pre - CNO), $p$=0.016 (CNO - washout), *n.s.* (pre - washout), Wilcoxon signed-rank test). Mean±SEM: 2.60±0.76, 8.18±2.94, 5.48±2.99 in pre, CNO and washout, respectively. For (D), (E) and (F), mean±SEM are indicated on the sides. *: $p<0.05$; **: $p<0.01$; ****: $p<0.0001$.

**Table S1. The total test time was not increased with Dimetindene maleate.** *p*-values according to one-tailed paired t-test.

| | Number of eyes | Placebo Mean±SD (sec) | Dimetindene maleate Mean±SD (sec) | p-value |
|---|---|---|---|---|
| **10-2 visual field test** | 16 | 321.63±37.18 | 324.13±49.30 | 0.7666 |
| **60-4 visual field test** | 16 | 419.75±81.20 | 425.50±84.89 | 0.7880 |

**Table S2. The retina's morphology was not altered by Dimetindene maleate.** See *Methods* for details on each measurement.

| | Number of eyes | Placebo Mean±SD | Dimetindene maleate Mean±SD |
|---|---|---|---|
| **Central macular thickness (µm)** | 16 | 262.50±12.98 | 262.68±13.19 |
| **RNFL thickness (µm)** | 16 | 104.56±3.48 | 104.37±2.91 |
| **Superficial plexus vascular density (%)** | 16 | 36.82±1.18 | 36.12±0.89 |
| **Deep plexus vascular density (%)** | 16 | 34.98±0.96 | 34.53±0.94 |
| **Choriocapillaris vascular density (%)** | 16 | 39.26±0.75 | 39.15±0.84 |
| **Choroidal vascular density (%)** | 16 | 37.28±0.59 | 37.25±0.46 |
